# The circadian clock is disrupted in pancreatic cancer

**DOI:** 10.1101/2022.11.01.514735

**Authors:** Patrick B. Schwartz, Manabu Nukaya, Mark E. Berres, Clifford D. Rubinstein, Gang Wu, John B. Hogenesch, Christopher A. Bradfield, Sean M. Ronnekleiv-Kelly

## Abstract

Disruption of the circadian clock is linked to cancer development and progression. Establishing this connection has proven beneficial for understanding cancer pathogenesis, determining prognosis, and uncovering novel therapeutic targets. However, barriers to characterizing the circadian clock in human pancreas and human pancreatic cancer – one of the deadliest malignancies – have hindered an appreciation of its role in this cancer. Here, we employed normalized coefficient of variation (nCV) and clock correlation analysis in human population-level data to determine the functioning of the circadian clock in pancreas cancer and adjacent normal tissue. We found a substantially attenuated clock in the pancreatic cancer tissue. Then we exploited our existing mouse pancreatic transcriptome data to perform an analysis of the human normal and pancreas cancer samples using a machine learning method, cyclic ordering by periodic structure (CYCLOPS). Through CYCLOPS ordering, we confirmed the nCV and clock correlation findings of an intact circadian clock in normal pancreas with robust cycling of several core clock genes. However, in pancreas cancer, there was a loss of rhythmicity of many core clock genes with an inability to effectively order the cancer samples, providing substantive evidence of a dysregulated clock. The implications of clock disruption were further assessed with a *Bmal1* knockout pancreas cancer model, which revealed that an arrhythmic clock caused accelerated cancer growth and worse survival, accompanied by chemoresistance and enrichment of key cancer-related pathways. These findings provide strong evidence for clock disruption in human pancreas cancer and demonstrate a link between circadian disruption and pancreas cancer progression.

**Author Summary:** The circadian clock is a regulator of human homeostasis. Dysfunction of the clock can lead to the development of diseases, including cancer. Although several cancers have been shown to have a dysfunctional clock which may alter prognosis or change treatment, this has been suggested but not demonstrated in pancreatic cancer. Investigation of this link is important because pancreatic cancer is highly lethal with few effective treatment options. Here we use recently pioneered bioinformatics approaches to assess clock functionality in human pancreatic cancer specimens, where we demonstrate that the clock is dysfunctional relative to normal pancreatic tissue. We then knocked out the core clock gene, *Bmal1*, in pancreatic cancer cells, which led to faster tumor growth and worse survival in mice and enhanced chemotherapeutic resistance to standard chemotherapy agents used in the treatment of pancreatic cancer. Collectively, our findings establish human pancreatic cancer as having clock dysfunction and clock dysfunction causing a more aggressive cancer.

## Introduction

The circadian clock is a conserved molecular feedback loop that regulates many signaling pathways to control metabolism, immunity, apoptosis, and other critical cellular functions in the body (1). At its core, the positive arm of the clock (*i.e.,* CLOCK and BMAL1 [also known as ARNTL or MOP3]) drives transcription of the negative arm, including PER1-3 and CRY1-2 (2,3). The negative arm represses the transcriptional activation of CLOCK and BMAL1, and a second interlocked loop involves the nuclear receptors RORα/β/λ and NR1D1-2 (also known as REV-ERBα/β), which activate and suppress BMAL1 expression, respectively. These, along with other core clock components, form a tightly regulated series of transcriptional-translational feedback loops that ensure rhythmic expression over 24 hours and function to maintain cellular and organ homeostasis (2–4).

Environmental cues influence the synchronization of circadian rhythms in various organ systems, and misalignment of external cues with the internal clock (*e.g*., shift work) can cause clock dysfunction with consequent metabolic derangements and pathologic states (4–6). For instance, circadian dysregulation has been strongly linked to obesity and diabetes, both risk factors for cancer (7–11). Concordantly, landmark studies have shown that disruption of the endogenous clock through mutations in or suppression of the core clock genes is intricately linked to tumor growth in several cancers (12–14). For example, knockout of *Bmal1* in *Kras*- and *p53*-mutant lung cancer causes marked tumor progression *in vivo* (13), while MYC-induced repression of *BMAL1* in human neuroblastoma drives decreased overall survival in a BMAL1-dependent manner (15). Importantly, targeting dysfunctional clock components in certain cancers has proven an effective treatment strategy (15,16). Thus, identifying an aberrantly functioning circadian clock in cancer can lead to key advancements such as understanding pathogenesis, prognosis, and uncovering novel therapeutic targets.

Although indeterminate, there is some evidence that the clock may be dysregulated in pancreatic ductal adenocarcinoma (PDAC), leading to a worse prognosis (17,18); this is alarming for a deadly malignancy where only 11% of patients survive beyond 5 years (19). To advance our understanding of how clock disruption impacts PDAC pathogenesis, and ultimately foster the identification of therapeutic targets, an essential first step is to establish that clock dysfunction exists in human PDAC. Unfortunately, to date, the cumulative data does not definitively support this assertion and is inconclusive. Prior studies have relied on contrasting expression differences between the core clock genes in tumor compared to normal pancreas as a basis for clock disruption (17,18,20,21). But differential expression alone is limited and does not provide insight into critical components of clock health such as relative amplitude, rhythmicity, or correlation of expression amongst core clock genes (22). For example, in the pancreas, phase advancement as a result of chronic jetlag causes differential expression of core clock genes despite maintaining a robust and healthy clock (*i.e.,* nearly identical relative amplitude, rhythmicity, and clock correlation) (23). Furthermore, in population-level data where sample time acquisition is unknown and the internal clock from individual patient tumors may not be in phase, the interpretation of differential expression for evaluating clock function becomes challenging. Thus, much more substantive data is required to affirm clock disruption in PDAC.

In pre-clinical studies (*e.g.*, mouse or cell-line models), the determinants of clock health can be identified by obtaining longitudinal data under controlled conditions (23,24). However, in human studies – particularly human pancreas – the requisite periodic data by multiple sampling is not feasible (or ethical). Therefore, alternative means of determining rhythmically expressed genes in humans are necessary.

Following the emergence of several bioinformatics tools, the principal aspects of clock health including relative amplitude, core clock gene correlation, and statistical determination of rhythmicity can be resolved when using the appropriate reference data (22,25–28). We recently generated a robust pancreas dataset demonstrating diurnally expressed genes over 48 hours (23). With this foundation, we were able to apply normalized coefficient of variation (nCV), clock correlation analysis, and cyclic ordering by periodic structure (CYCLOPS) to evaluate the principal aspects of clock function, respectively, and test the hypothesis that the circadian clock is disrupted in human PDAC (22,25,28). While on the surface this may be construed as a simplistic hypothesis, several limitations have hindered the ability to evaluate the circadian clock in human pancreas, including the lack of human periodic data and an absence of a reference transcriptional dataset.

Here, we employed nCV, clock correlation analysis, and CYCLOPS on publicly available human population-level expression data from The Cancer Genome Atlas (TCGA) and Clinical Proteomic Tumor Analysis Consortium 3 (CPTAC-3) datasets to determine the health of the circadian clock in PDAC and adjacent normal tissue (29,30). We identified an intact circadian clock in human normal pancreas with robust cycling of several core clock genes. We also found a markedly weakened clock in the cancer tissue, providing substantive evidence of a dysregulated clock in PDAC. These findings represent significant advancements in evaluating clock function in PDAC, and the potential clinical implications of clock disruption were further assessed with a pre-clinical PDAC model. We used CRISPR/Cas9 technology to selectively target *Bmal1* and examined the effects of clock dysfunction *in vitro* and *in vivo*. This revealed that loss of clock function caused accelerated cancer growth, diminished survival, enrichment of key cancer-related pathways, and resistance to commonly used cytotoxic chemotherapies for PDAC. These findings provide strong evidence for circadian clock disruption in human PDAC and demonstrate a link between circadian disruption and pancreas cancer progression.

## Results

### The Clock in Human Pancreatic Ductal Adenocarcinoma is Less Robust Than in Normal Pancreatic Tissue

To test the hypothesis that the circadian clock is disrupted in human pancreatic ductal adenocarcinoma (PDAC), we used our bioinformatics pipeline that consists of clock correlation, normalized coefficient of variation (nCV), and cyclic ordering by periodic structure (CYCLOPS) (**Fig 1**). We performed an analysis on publicly available The Cancer Genome Atlas (TCGA) and Clinical Proteomics Tumor Analysis Consortium 3 (CPTAC-3) pancreatic datasets (29,30). However, we first needed to ensure the validity of the pipeline in the pancreas since these had not been previously applied to pancreatic tissue. We assessed the clock in our existing murine normal circadian and chronic jetlag pancreas RNA-sequencing (RNA-seq) datasets by examining the clock correlation matrix and nCV (23). These datasets contained pancreas samples acquired every 4 hours for 48 hours in male (n = 3 at each timepoint for each condition) and female (n = 3 at each timepoint for each condition) mice under standard lighting (normal circadian, n = 72) and chronic jetlag (n = 72) conditions. Chronic jetlag is known to affect the phase of gene expression but not the relative amplitude (rAMP) or the correlation of the core clock genes in the pancreas (23).

**Fig 1:**
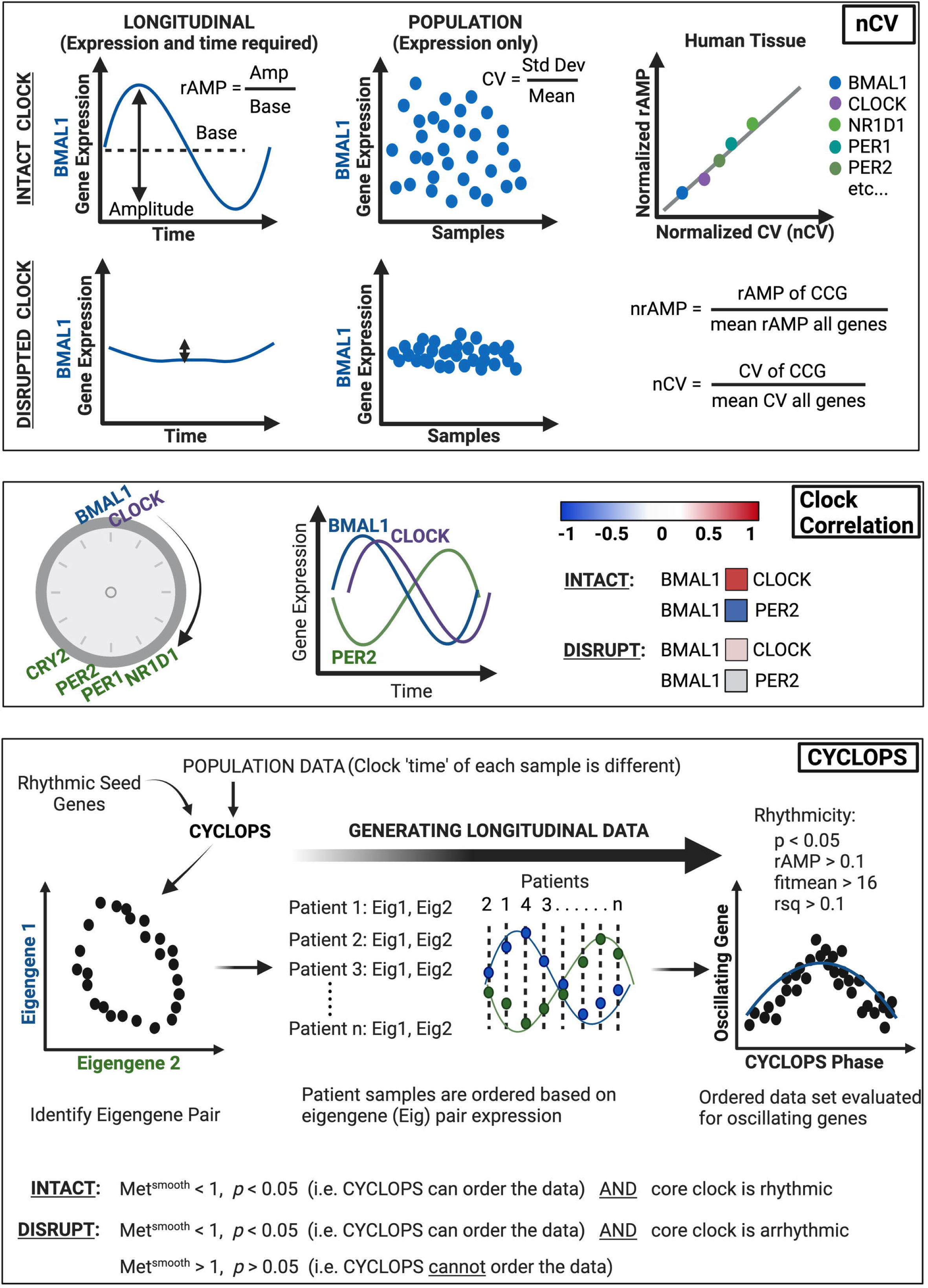
Bioinformatics pipeline for circadian function. Recently, there have been three separate bioinformatics tools that have substantially improved the ability to determine circadian clock function in population-level transcriptional data (22,25–28). **<u>Normalized coefficient of variation (nCV)</u>**: Clock gene expression produces robust oscillations with the amplitude of the oscillation defined by the difference between peak and trough, and relative amplitude (rAMP) determined by the ratio between amplitude and baseline level of expression (<u>upper left</u>). Clock disruption (such as with clock gene knockout) causes suppressed rAMP, and thus rAMP can be considered a strong determinant of clock function (<u>lower left</u>). Because rAMP is calculated from time-labeled samples, it can only be measured in human population data when sample time acquisition is known (*e.g.,* MetaCycle) or when using programs to predict longitudinal time-course data (*e.g.,* CYCLOPS). To overcome this limitation and assess core clock gene ‘robustness’ of oscillation in human data (without labeled time), normalized coefficient of variation (nCV) is used. The nCV is the coefficient of variation (CV) of the core clock gene (in the population data) divided by the mean CV of all genes in the dataset. And the CV is determined by the standard deviation of the gene expression divided by its average expression. Thus, when the circadian clock is intact, circadian genes in large population data sets would be expected to have a relatively high nCV due to oscillating gene expression (<u>upper middle</u>). Conversely, with a disrupted clock, the variation of the core clock genes would be diminished (oscillation is suppressed), resulting in a relatively low nCV (<u>lower middle</u>). Importantly, the nCV was found – across multiple cancer types and tissues – to correspond directly to the normalized rAMP (<u>upper right</u>) (25); it can thus be used confidently as a surrogate for rAMP, which is a key determinant of clock health. Of note, the rAMP (and thus nCV) of core clock genes differs due to differences in amplitude, and so the between tissue difference (*e.g.,* normal vs cancer) for each core clock gene is the value of importance. **<u>Clock Correlation</u>**: When the circadian clock is intact, there is an expected progression of core clock gene expression, where the positive arm of the clock (*e.g*., *BMAL1*, *CLOCK*) drives transcription of the negative arm of the clock (*e.g., PER*, *CRY*). Thus, when *BMAL1* expression peaks, *CLOCK* expression should also be near its peak, and this should be evident in population-level data (*i.e.,* can look at the spread of individual expression points across the population). In turn, when *BMAL1* expression peaks, negative arm members (e.g., *PER1-2*) should be at a trough. Concordantly, strong positive correlations (0.5 < ρ < 1, red) should be apparent among transcriptional activators (e.g., *BMAL1* and *CLOCK*) and among transcriptional repressors (e.g., *NR1D1* and *PER2*), and a strong negative correlation (-0.5 > ρ > -1, blue) should be present amongst activators and repressor targets (e.g., *BMAL1* and *PER2*). If this is the case, then the clock is intact; if these correlations are not preserved (*i.e.,* ρ ∼ 0), this indicates the clock is disrupted. Using the set of core clock genes and clock- associated genes, the correlation of each gene against the others is determined by Spearman’s rho (ρ) and mapped in matrix form. This experimental matrix is then compared to a baseline correlation matrix from the mouse circadian gene atlas using the Mantel test, which compares the correlation between the two matrices to produce a z-statistic (z-stat). A higher z-stat value corresponds to a correlation matrix that is closer to the circadian gene atlas baseline (*i.e.,* highly preserved clock correlation). **<u>CYCLOPS</u>:** Intrinsic to circadian clock genes is rhythmic expression. CYCLOPS (cyclic ordering by periodic structure) is an algorithm that can identify rhythmic (longitudinal) data from population data where sample time acquisition is unknown. The basis for this type of program is that each sample has a different ‘clock time’ due to differences in time of sample acquisition and environmental factors such as sleep/wake cycle or shift worker status (*e.g.,* 15-20% of patients are night shift workers). Seed genes that are known to be rhythmic in the tissue of interest (*i.e.,* pancreas) are inputted into CYCLOPS, and the population level data is reduced to two vectors (Eigengene 1 and Eigengene 2) derived from the seed genes. When plotted, the optimal Eigengene pair will demonstrate an ellipse, which indicates that the two Eigengenes are rhythmic and anti-phasic – this pair can then be used for each patient sample to determine the sample ‘time’, and thus the order of that sample relative to the 24-hr period (phase). By incorporating enough patients, population-level data can be transformed into longitudinal data. Subsequently, with the patient sample dataset ordered by CYCLOPS (by the Eigengene pair), individual genes (*e.g.,* oscillating genes or core clock genes) can be evaluated for rhythmicity. An intact circadian clock is indicated by the ability of CYCLOPS to order the data (statistically) and the identification of rhythmically expressed core clock genes. Meanwhile, a disrupted clock is designated as arrhythmic core clock gene expression or an inability of CYCLOPS to order the data (statistically). Statistically significant rhythmic gene expression is determined by *p* < 0.05, rAMP > 0.1, fitmean > 16 and goodness of fit (rsq) > 0.1. The fitmean value can be conceptualized as a mean level of expression of that gene across the dataset, and therefore a minimum level of expression (fitmean > 16) is required for rhythmicity cutoff. The goodness of fit of the experimental values by cosinor regression is calculated by R squared (rsq). CYCLOPS reordering is assessed by the Met^smooth^ and Stat^err^ (significant reordering: Met^smooth^ < 1 and Stat^err^ < 0.05). The Met^smooth^ compares the smoothness of the reconstructed circular trajectory versus a linear ordering based on the first principal component, while the Stat^err^ (we designate as *p*-value) is analogous to the F statistic of a typical nested regression and compares the model fit when a circular rather than linear bottleneck node is used. A full mathematical derivation has been previously outlined by Anafi *et al.* (28) Figure created with BioRender.com.

Correspondingly, we found that the core clock relationships (as measured by the z-statistic or standard deviations above/below population mean) remained intact on the correlation matrix when compared to the baseline clock correlation matrix for both normal and chronic jetlag conditions (*p* < 0.001, z-statistic = 54.96 vs. *p* < 0.001, z-statistic = 54.08) (**Fig 2A**) (31,32). Further, the nCV of 11 clock genes remained unchanged (*p* = 0.76) between the normal circadian (mean nCV (± standard error) = 1.78 (± 0.27)) and chronic jetlag (mean nCV = 1.9 (± 0.28)) conditions (**Fig 2B-C**) – consistent with the murine pancreatic clock being strongly rhythmic and intact, matching precisely what we had found in our prior longitudinal analysis (23).

**Fig 2:**
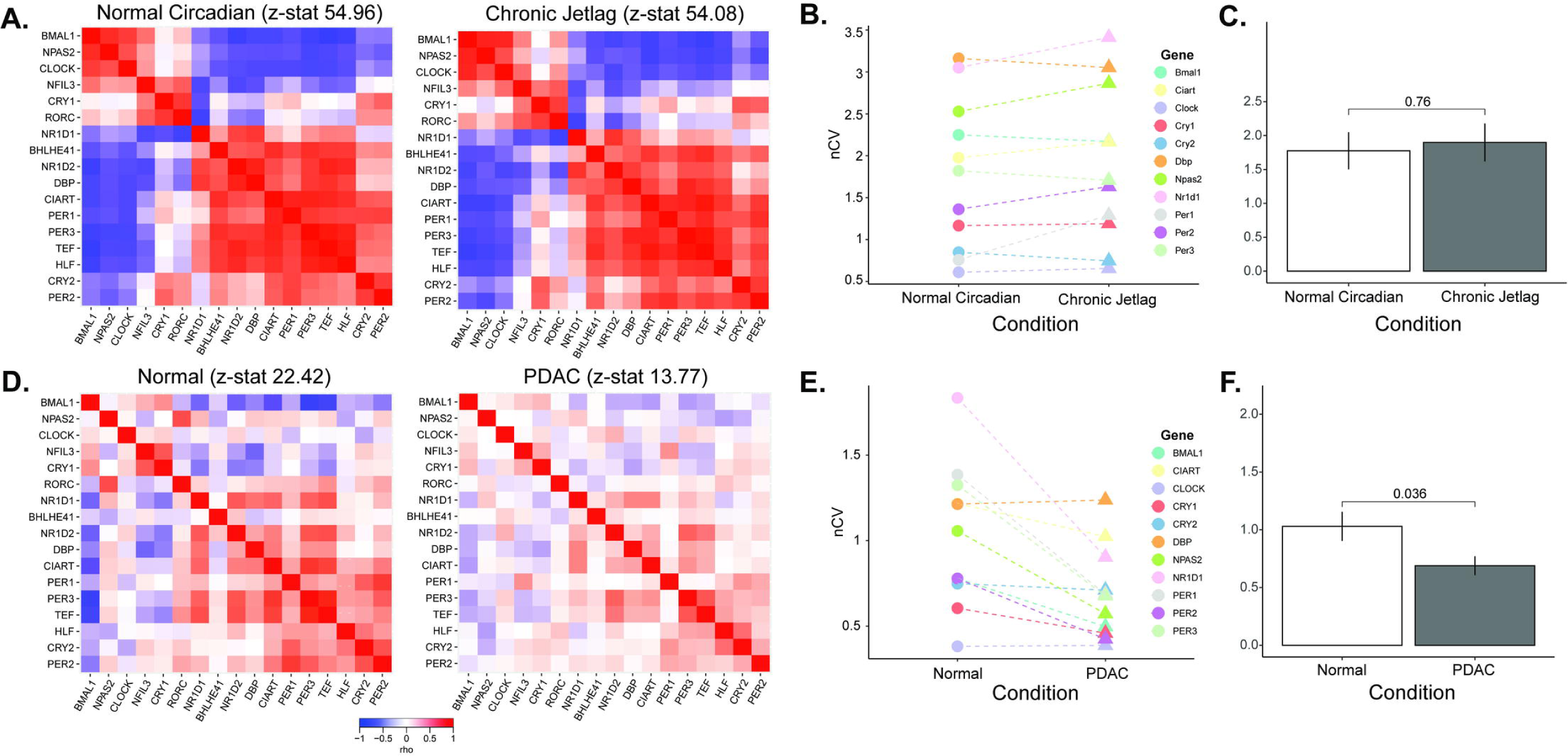
The human pancreatic clock robustness is diminished after malignant transformation. **A**. Clock correlation matrices were created for normal circadian (left, n = 72) and chronic jetlag (right, n = 72) mouse pancreas samples. **B, C**. The normalized coefficient of variation (nCV) was calculated to determine the robustness of the pancreatic clock for the normal circadian (circles) and chronic jetlag (triangles) pancreas samples, with the mean nCV in each group also depicted. Each color indicates a different gene evaluated. **D.** Clock correlation matrices were created for normal (left; n = 50) and pancreatic ductal adenocarcinoma (right; n = 318) samples. **E, F**. The normalized coefficient of variation (nCV) was calculated to determine the robustness of the pancreatic clock for the normal (circles) and PDAC (triangles) samples, with the mean nCV in each group also depicted. Each color indicates a different gene evaluated.

CYCLOPS was then used to reorder the mouse pancreas samples by their predicted phase, and cluster reordering of samples by CYCLOPS was analyzed for appropriate phase progression compared to the known zeitgeber time (ZT) of sample collection. CYCLOPS reordering can be assessed by the Met^smooth^ and Stat^err^ (significant reordering: Met^smooth^ < 1 and Stat^err^ < 0.05). A full mathematical derivation has been previously outlined by Anafi *et al* (28). Notably, CYCLOPS accurately (phase appropriately) reordered both the normal (*p* < 0.01; Met^smooth^ = 0.90) and the chronic jetlag conditions (*p* < 0.01; Met^smooth^ = 0.97), validating the pipeline for use in the pancreas (**Supplemental Fig 1A-C**). Collectively, these results confirmed the ability of nCV, correlation matrix, and CYCLOPS to determine the robustness of the circadian clock in the pancreas.

Considerable data support the tumor suppressor role of the circadian clock and the assertion that circadian disruption is present in human cancer (33). In human PDAC, the supposition is that the clock may be altered, but supporting data is limited to the relative expression of single genes and is consequently inconclusive (18,21,34). We therefore applied our pipeline to determine clock health in PDAC using the TCGA and CPTAC-3 pancreatic datasets (22,26–30). After processing and batch correction of the matched 50 available normal and 318 available PDAC samples (**Supplemental Fig 2**), we examined the correlation matrix between the core clock genes (CCGs) and found that there was a weaker relationship amongst the CCGs in PDAC compared to normal (z-statistic PDAC = 13.77 vs. z-statistic normal = 22.42) (**Fig 2D**). Concordantly, we found that there was a clear decrease (*p* = 0.04) in the nCV between PDAC (mean nCV = 0.69 (± 0.08)) and normal (mean nCV = 1.03 (± 0.13)) (**Fig 2E-F**), indicating a weaker clock in PDAC compared to normal pancreas. We then applied our pancreatic CYCLOPS analysis which significantly reordered the normal samples (*p* < 0.01; Met^smooth^ = 0.99), indicating an intact clock. This was further reinforced when evaluating the predicted phase of CCGs and clock-controlled genes using CYCLOPS. We found that the phase sequence of normal pancreatic samples was similar to the phase sequence of mouse pancreatic samples (**Fig 3A-B**), with the phase of expression in the human genes equivalent to the mouse genes in 12 out of 17 CCGs and clock-controlled genes and ROR-phased genes largely peaking before E-box phase genes. Based on the CYCLOPS ordering, we analyzed the proportion of genes that were rhythmically expressed in normal pancreas. We found that 4,034/18,196 (22.17%) of normal pancreatic genes were rhythmic, (**Supplemental Data File 1**). We found that several key CCGs and clock-controlled genes were rhythmically expressed in normal samples on cosinor analysis, including *CLOCK*, *PER1*, *PER3*, *NR1D2*, *RORC*, *NFIL3, TEF*, *NR1D1, BHLHE40,* and *NPAS2* (**Fig 4A and Supplemental Table 1**). The CCGs *BMAL1* (*p* = 0.07, rAMP = 0.8) and *CRY1* (*p* = 0.08, rAMP = 0.48) were nearly rhythmic in the normal samples based on the pre-defined cutoff. Finally, phase set enrichment analysis (PSEA) was used to determine phase-dependent gene enrichment. The top 15 significant pathways were selected for analysis (**Fig 4B**). Consistent with our prior murine pancreas data, we found that normal pancreas was associated with time-dependent metabolic gene pathway enrichment (23,35). In aggregate, this data confirms the health of the circadian clock in tumor-adjacent normal pancreatic tissue, which was expected but not previously shown to our knowledge.

**Fig 3:**
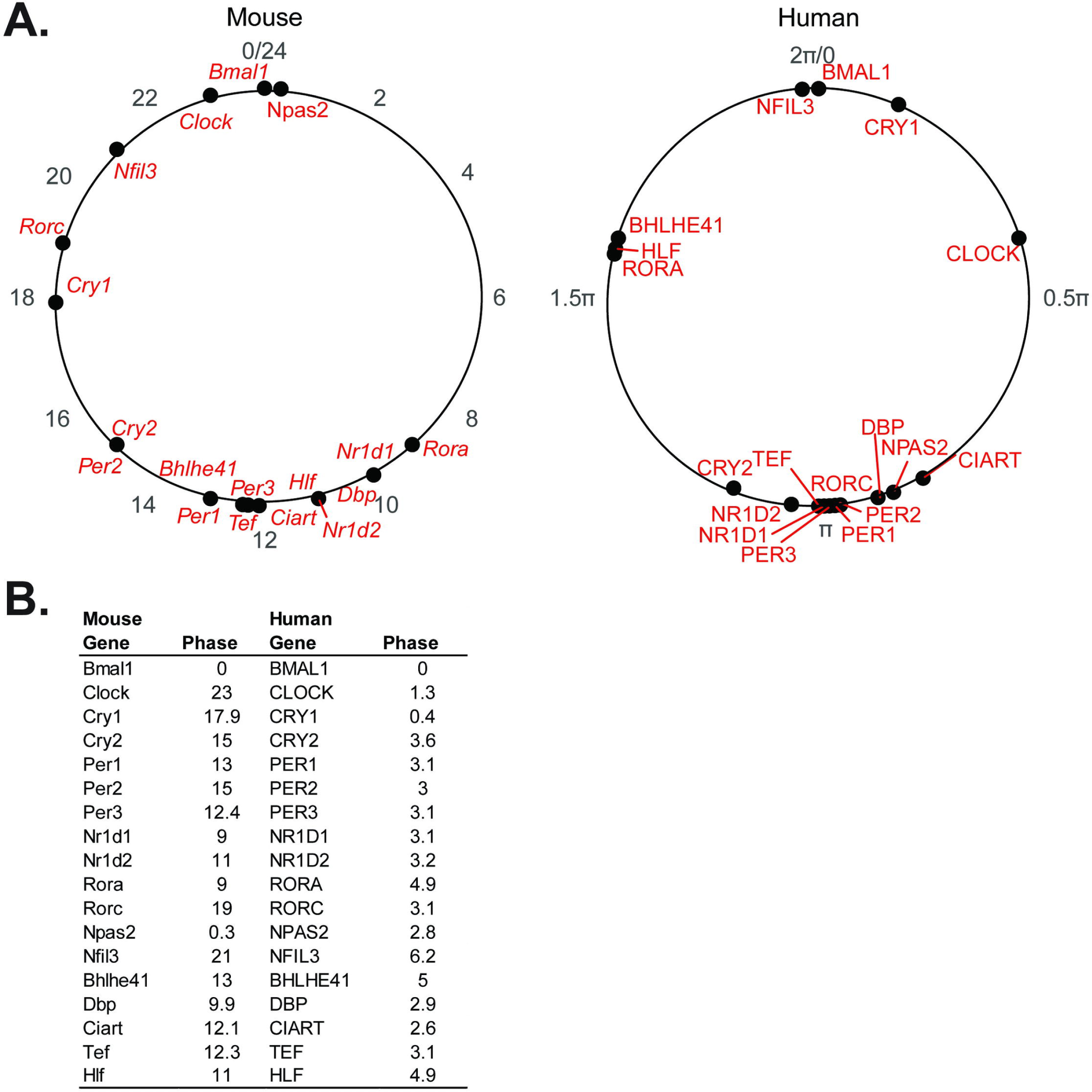
Clock phase predictions demonstrate conserved clock relationships: **A.** Graphical representation and **B.** table of the CCG and clock-controlled gene phase prediction from the cosinor analysis of CYCLOPS reordered human pancreas samples (0 to 2π; n = 50) and Metacycle predicted phases of mouse pancreas samples (0 to 24; n = 72), relative to *BMAL1* (human) and *Bmal1* (mouse) phase.

**Fig 4:**
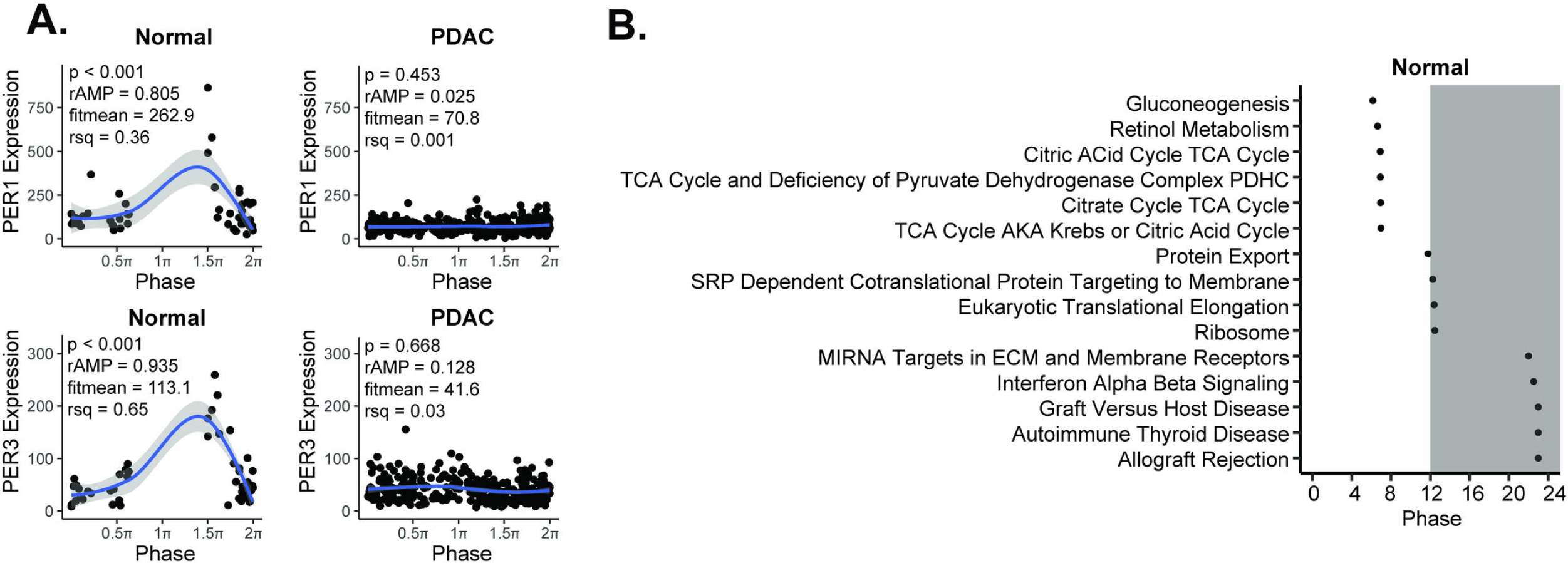
The human pancreatic cancer tumor clock is dysfunctional relative to the normal pancreas. **A**. CYCLOPS was used to reorder samples from normal (n = 50) and pancreatic ductal adenocarcinoma (n = 318) TCGA and CPTAC-3 samples. Each point represents an individual patient with order determined by paired eigengenes. Plots from several core clock genes reordered by CYCLOPS in normal tissue as compared to best reordering in PDAC (**A**) are shown – ordered from 0 to 2π. Shading around the blue regression line indicates the 95% confidence interval. Relative amplitude (rAMP) for each gene is determined by dividing the amplitude by the fitted expression baseline (fitmean). The fitmean value can be conceptualized as a mean level of expression of that gene across the dataset, and therefore a minimum level of expression (fitmean > 16) is required for rhythmicity cutoff. The goodness of fit of the experimental values by cosinor regression was calculated by R^2^(rsq). Using these collective parameters, a *p* < 0.05, rAMP > 0.1, goodness of fit (rsq) > 0.1, and fitmean > 16 were considered rhythmic (27). **B**. Graphical representation of Phase Set Enrichment Analysis (PSEA) of normal pancreatic samples, ordered by phase of expression.

Despite the validation steps and success in ordering normal pancreas, CYCLOPS **could not** reorder PDAC samples (*p* = 0.43; Met^smooth^ = 0.99), revealing a less robust (or disrupted) clock in PDAC compared to normal. This inability to reorder strongly supports the attenuated circadian clock in PDAC identified with nCV and clock correlation analysis. We selected the best reordering available based on the eigengenes to enable visual representation of the PDAC samples compared to normal pancreas samples. This also enabled us to estimate as best as possible the rAMP of the PDAC samples, to coincide with the visual representation. Consistent with the nCV analysis, we found that rAMP decreased substantially in every core clock gene from normal to tumor (**Supplemental Table 1**), as evidenced by *PER1* and *PER3* expression (**Fig 4A**). Collectively, the weaker clock correlation, markedly reduced nCV, and CYCLOPS analysis of human population-level data show that the circadian clock is significantly disrupted in PDAC.

### Creation of a Circadian Dysfunction Pancreatic Cancer Model

After demonstrating that the clock in human PDAC was dysfunctional, we sought to understand the potential clinical implications of clock disruption. Identifying an aberrant clock in other cancers has consistently demonstrated accelerated cancer progression and worse prognosis, while simultaneously revealing novel therapeutic targets (13–16). Moreover, there appears to be a putative correlation between suppressed *BMAL1* expression and poor prognosis in patients with PDAC (17,18). Therefore, we hypothesized that disruption of the circadian clock in PDAC would cause accelerated cancer progression. To test this hypothesis, we used a syngeneic *Kras*- and *Trp53*-mutant pancreas cancer cell line (KPC) and employed CRISPR/Cas9 technology to functionally knockout *Bmal1* (KPC-BKO) so that we could examine the effects of clock dysfunction *in vitro* and *in vivo* (36). The reason for mutating *Bmal1* in the PDAC cells was several-fold: i) KPC cells demonstrated an intact clock (**Fig 5D**) necessitating core clock gene modulation, ii) *BMAL1* is a central transcriptional regulator of the circadian clock machinery and functional knockout of this single gene abolishes circadian function (37,38), iii) suppressed *BMAL1* expression has been found in several human cancers, correlating with worse prognosis (13,15,17,18,20,21,39), iv) *BMAL1* expression is decreased in tumor compared to normal tissue in human PDAC (18), v) we similarly found a substantially dampened nCV (rAMP) of *BMAL1* expression in PDAC compared to normal tissue (*PER1*, *PER3*, *NR1D1,* and *BMAL1* displayed the greatest decrease in nCV), and vi) segregating patients with PDAC into high and low *BMAL1* expression appears to be prognostic for survival outcomes (17,18,21).

**Fig 5:**
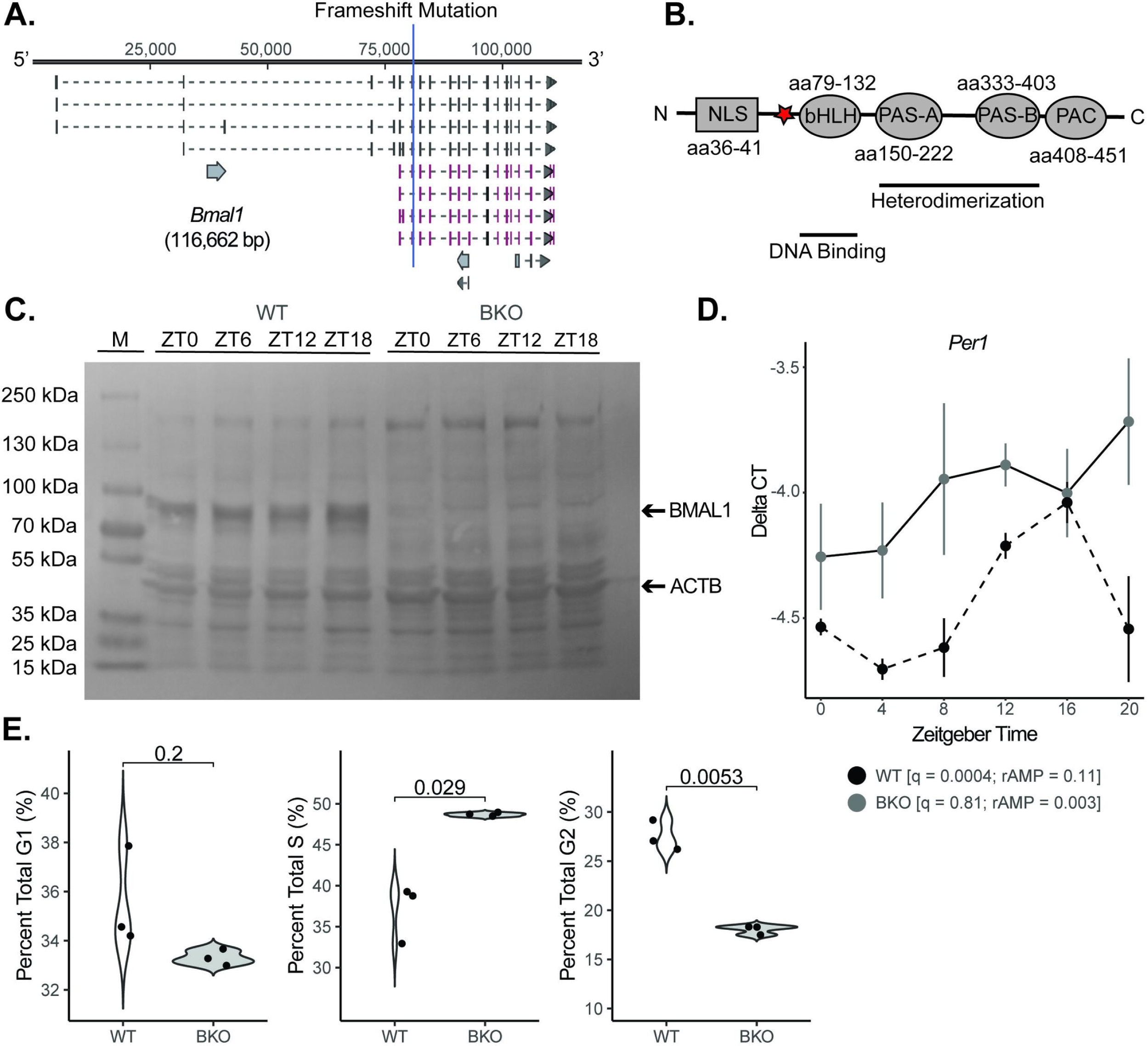
Creation of a mouse clock dysfunction cell line model. **A**. The core clock gene *Bmal1* was mutated (functional knockout) in KPC murine pancreatic cells with CRISPR-Cas9 genome editing technology (BKO) with the frameshift site indicated by the blue line. **B.** The frameshift mutation was induced upstream of the basic helix loop helix (bHLH) domain (red star). The resulting protein lacked critical downstream elements for functionality, including the PAS-A, PAS-B, and PAC domains. **C.** Western blot demonstrating the loss of BMAL1 protein across 24 hours for wild-type (WT) and BKO cells (M = marker lane). **D.** The mean (± standard error) *Per1* mRNA expression of synchronized KPC (n = 3 at each time point; black dashed) and KPC-BKO (n = 3 at each time point; grey) cells from ZT0-ZT24. **E.** Graphs comparing the percent of cells in G1, S, and G2 in WT (n = 3; white) and BKO (n = 3; grey) KPC cells.

Consequently, guide RNAs were designed to target exon 8 of the *Bmal1* gene, which resulted in the insertion of adenine on one allele and a 2 base pair deletion on the other between nucleotide 81,074 and 81,075 at amino acid 73 (GRCm38) (**Fig 5A**). The result of both mutations was a frameshift just upstream of the basic helix loop helix (bHLH) domain known to assist the PAS A domain in heterodimerization with its binding partner CLOCK (**Fig 5B**) (40,41). Confirmatory western blotting revealed the presence of protein in wild-type (WT) cells and an absence of protein in *Bmal1* knockout (BKO) KPC cells (**Fig 5C**). To evaluate clock functionality, mRNA was isolated at 4-hour intervals (n = 3 per condition) for 24 hours after cell synchronization, and we performed RT-qPCR for *Per1*, a core downstream repressor of the positive arm of the molecular clock and a gene that demonstrated rhythmicity in our human normal CYCLOPS analysis (**Fig 5D**). Rhythmicity was assessed with the meta2d function of the Metacycle package (42). KPC (WT) cells exhibited an intact and robust circadian clock with a *q* = 0.0004 and rAMP = 0.11 for *Per1*. Although still rhythmic, when compared to normal murine pancreas core clock gene function from our prior longitudinal transcriptional dataset (normal pancreas *Per1*: *q* = 0.0001, rAMP = 0.394), the relative amplitude was diminished (23). To further evaluate the functionality of the clock in KPC cells, luciferase activity driven by the *Per2* promoter was measured. Luciferase activity in two separate clones was highly rhythmic (Clone 1: *q* = 5.51E-5, rAMP = 0.44; Clone 2: *q* = 1.95E-7, rAMP = 0.39) (**Supplemental Fig 3**). Consistent with the comparison of *Per1* expression in KPC versus our prior normal murine pancreas dataset, the relative amplitude of *Per2* was diminished in the KPC lines compared to normal pancreas tissue (*q* = 1.39E-7, rAMP = 0.82). Thus, although rAMP was diminished in the murine pancreas cancer cell line compared to normal pancreas tissue, clock function was still intact. Conversely, KPC-BKO cells exhibited no detectable clock function with a *q* = 0.81 and rAMP = 0.003 for *Per1* expression (**Fig 5D)**. In addition, we looked for differences in the cell cycle consequent to clock dysfunction as the two pathways are known to interact (43,44). For instance, Bieler *et al.* identified robust synchrony between cell division and circadian cycle in NIH3T3 fibroblasts that contained a *Rev-Erba*- YFP reporter (43), while Feillet *et al*. used a similar system (NIH3T3 fibroblasts) to identify a coordinate entrainment in unsynchronized cells as well as cells that were dexamethasone synchronized (44).

Consistent with this known cross-talk between the circadian clock and cell cycle, KPC-BKO cells demonstrated alterations in the cell cycle compared to KPC cells (**Fig 5E**) (20). BKO cells were found to have a higher mean (± standard error) percentage of cells in S phase (48.66 (± 0.13) vs. 36.96 (± 2.04); *p* = 0.029) and a lower percentage in G2 (18.03 (± 0.27) vs. 27.49 (± 0.89); *p* = 0.0053). There were no differences between KPC and KPC-BKO cells in G1 (35.56 (± 1.19) vs. 33.31 (± 0.17); *p* = 0.2). Taken together, these data demonstrate the successful creation of a novel PDAC cell line with an abolished clock (*Bmal1* gene mutation), which can be used to elucidate the repercussions of clock disruption in PDAC.

### Clock Dysfunction Accelerates Pancreatic Cancer Growth

After establishing the model, we sought to understand how a dysfunctional clock impacted cell growth. KPC and KPC-BKO cells were grown in culture and injected into the right flanks of syngeneic C57BL/6J mice. Tumor growth was measured every 3-4 days for a total of 28 days to understand differences in primary tumor growth. We found that BKO caused an accelerated growth pattern, resulting in higher mean (± standard error) weight tumors at the study conclusion compared to KPC-derived tumors (438.02 ± 48.84 mg vs. 280.11 ± 42.73 mg; p = 0.02) (**Fig 6A-B**). These findings were evaluated and independently confirmed in a second identically created *Bmal1* mutant (functional knockout) clone (**Supplemental Fig 4A-C**). Perhaps more relevant to human PDAC – given the aggressiveness of this cancer – we assessed the effect of clock disruption on survival by implanting KPC and KPC-BKO heterotopic tumors and observing the mice until moribund status or lethality (**Fig 6C**). On Kaplan Meier log-rank analysis, mice harboring tumors derived from KPC-BKO cells experienced worse survival than mice bearing KPC-derived tumors (median survival: 52 versus 75 days, *p* = 0.04). Thus, clock dysfunction resulted in accelerated tumor growth *in vivo*, leading to worse overall survival.

**Fig 6:**
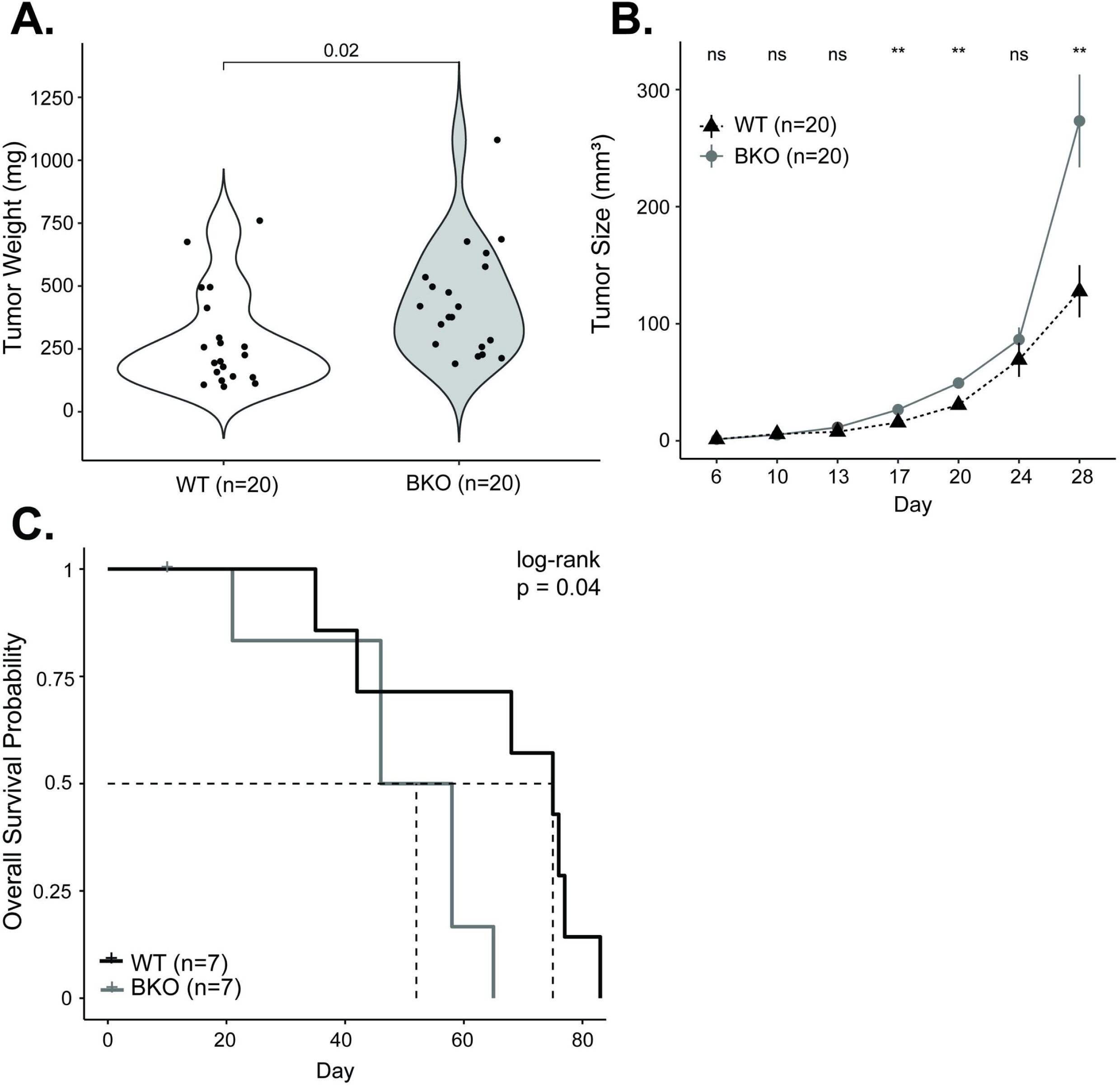
Mutation of Bmal1 promotes pancreatic cancer progression. **A.** Comparison of mouse tumor weight at 28 days for WT (n = 20; white) and BKO (n = 20) KPC tumors. **B.** WT (n = 20; black) and BKO (n = 20; grey) mean (± standard error) tumor size (in mm^3^) over the 28-day growth period. **C.** Kaplan Meier survival curve for mice implanted with WT (n = 7; black) and BKO (n =7; grey) KPC tumors. The dotted lines indicate the median survival. [ns = not significant, * = p < 0.05, ** = p < 0.01]

### Loss of Bmal1 Promotes the Enrichment of Cell Growth Pathways

To determine the possible etiologies of clock disruption causing accelerated PDAC progression, we compared the transcriptomic profiles of KPC and KPC-BKO cells. The principal component analysis revealed clear transcriptional profile differences between each condition (**Fig 7A**). Differential gene expression was quantified with edgeR and of the 15,110 genes, 5,235 (34.65%) were upregulated and 5,113 (33.84%) were downregulated in KPC-BKO compared to KPC cells (**Fig 7B, Supplemental Data File 2**) (45). When we examined the CCGs known to control the circadian cycle, we found that 12/15 examined clock genes were differentially expressed, including *Per1*, *Per2*, *Per3*, *Cry1*, *Cry2*, *Nr1d1*, *Nr1d2*, *Bhlhe40*, *Bhlhe41*, *Npas2*, *Arntl2*, and *Dbp* (All q < 0.05) (40). We then performed a KEGG pathway analysis to examine for enrichment of pathways as a result of *Bmal1* knockout in the PDAC cells (**Fig 7C**). We found that there was an enrichment of pathways important for cellular adhesion, such as ECM-Receptor Interaction, Cell Adhesion, and Focal Adhesion, as well as several cellular growth pathways including PI3K-AKT Signaling Pathway, MAPK Signaling Pathway, and Rap1 Signaling Pathway. Collectively, these data indicate that clock disruption in the KPC cells results in significant core clock gene changes and a transcriptional shift that alters key cancer promotion-related pathways such as cellular attachment and proliferation.

**Fig 7:**
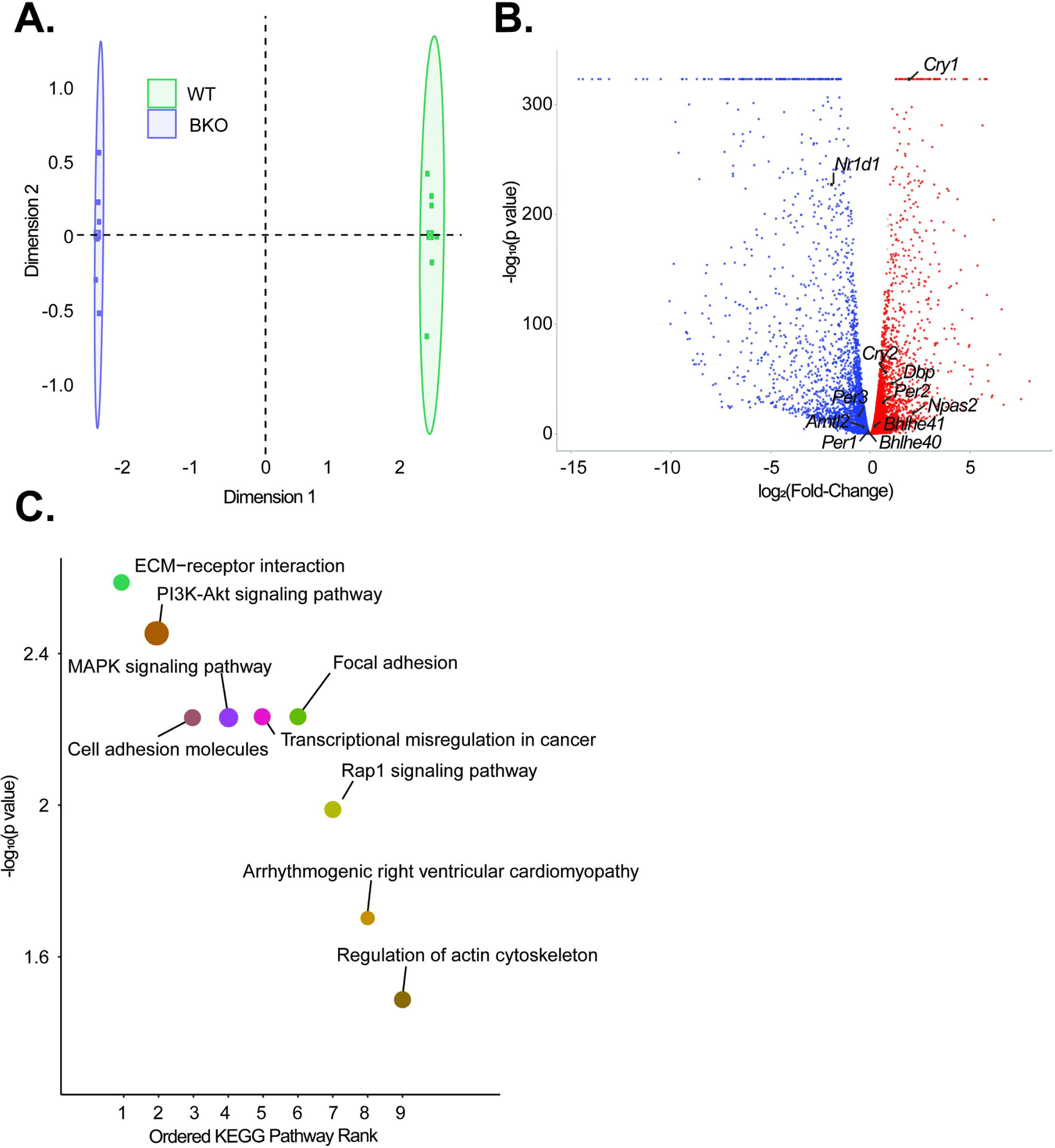
Functional knockout of Bmal1 promotes widespread transcriptomic alterations and activation of multiple oncogenic pathways. **A.** Principal Component Analysis (PCA) demonstrating overall differences in expression between WT (green) and BKO (purple) samples (n = 6 each) **B.** Volcano plot showing genes that were significantly upregulated (red) and downregulated (blue) on differential gene expression (DGE) analysis (q < 0.05). All significant core clock genes are highlighted **C.** KEGG analysis was then performed, and the top 9 pathways ordered by significance are shown. For each pathway, the size of each dot corresponds to the number of genes involved in each pathway.

### Cell Survival is Increased with the Loss of Bmal1 through Alterations in Multiple Cell Death Pathways

A hallmark of cancer progression is the inhibition of apoptosis (46). This phenotype is readily apparent in response to chemotherapy. In particular, the mechanism of action of gemcitabine (inhibition of DNA synthesis) and paclitaxel (microtubule stabilization) – backbones in PDAC therapy – is to ultimately induce apoptosis (47,48). To understand the potential implications of clock disruption for PDAC patients undergoing treatment, we subjected KPC and KPC-BKO cells to 72 hours of chemotherapeutic treatment with either gemcitabine or paclitaxel (49). We found that the activity of cell death pathways through apoptosis (measured by Caspase 3/7 activity) was blunted in response to both chemotherapeutic agents as a result of clock dysfunction (**Fig 8A-B**). We also assessed cytotoxic cell death, as measured by Dead-Cell Protease activity, and found that clock disruption promoted resistance to cytotoxic cell death induced by gemcitabine and paclitaxel (**Fig 8C-D**). These findings were also confirmed in a separately evaluated independent *Bmal1*-mutant clone (**Supplemental Fig 4 D-G**). Prior work has implicated alteration of Trp53 signaling to impact cell survival by suppressed apoptosis (20), but KPC cells (both WT and BKO) are *Trp53* mutant (*Trp53^R172H^*) indicating alternative mechanisms of heightened resistance in the KPC- BKO cells. Gemcitabine resistance in PDAC often occurs due to the downregulation of the channel protein hENT1 (*SLC29A1* gene), or deoxycytidine kinase (*DCK* gene) which activates gemcitabine once inside the cell (50). However, these were only marginally downregulated (1.05-fold, *q* = 0.04 and 1.08- fold, *q* = 0.02) due to *Bmal1* knockout, and thus unlikely to contribute to the differences seen with clock disruption (**Fig 9A**). Furthermore, these were arrhythmic based on rhythmicity cut-off values (*p* < 0.05, rAMP > 0.1, goodness of fit (rsq) > 0.1, and fitmean > 16) when examining our human samples, suggesting a lack of circadian control (**Fig 9B**, **Supplemental Table 1**). Meanwhile, paclitaxel drug resistance is thought to occur mostly through upregulation of drug efflux transporter proteins (P-glycoprotein also known as multidrug-resistance associated protein), but these ATP-binding cassette transporter proteins (*Abcb1a and Abcb1b* genes) were instead significantly *downregulated* (8.2-fold, *q* < 0.0001 and 2.1-fold, *q* < 0.0001) in KPC-BKO vs KPC cells (**Fig 9C**) (48). Concordantly, the *ABCB1* gene was rhythmic with elevated expression in the human normal samples (*p* = 0.015, rAMP = 0.95, fitmean = 133, rsq = 0.57) compared to the human PDAC samples that showed low levels of expression (mean expression = 22) (**Fig 9D**, **Supplemental Table 1**). Thus, using this combined data analysis, we found that the commonly described resistance mechanisms for gemcitabine and paclitaxel were not identified, underscoring the complexity of clock disruption induced chemotherapeutic resistance in cancer. Regardless, these data demonstrate that clock dysfunction promotes broad resistance in PDAC including multiple cell death pathways in the setting of two different PDAC backbone agents.

**Fig 8:**
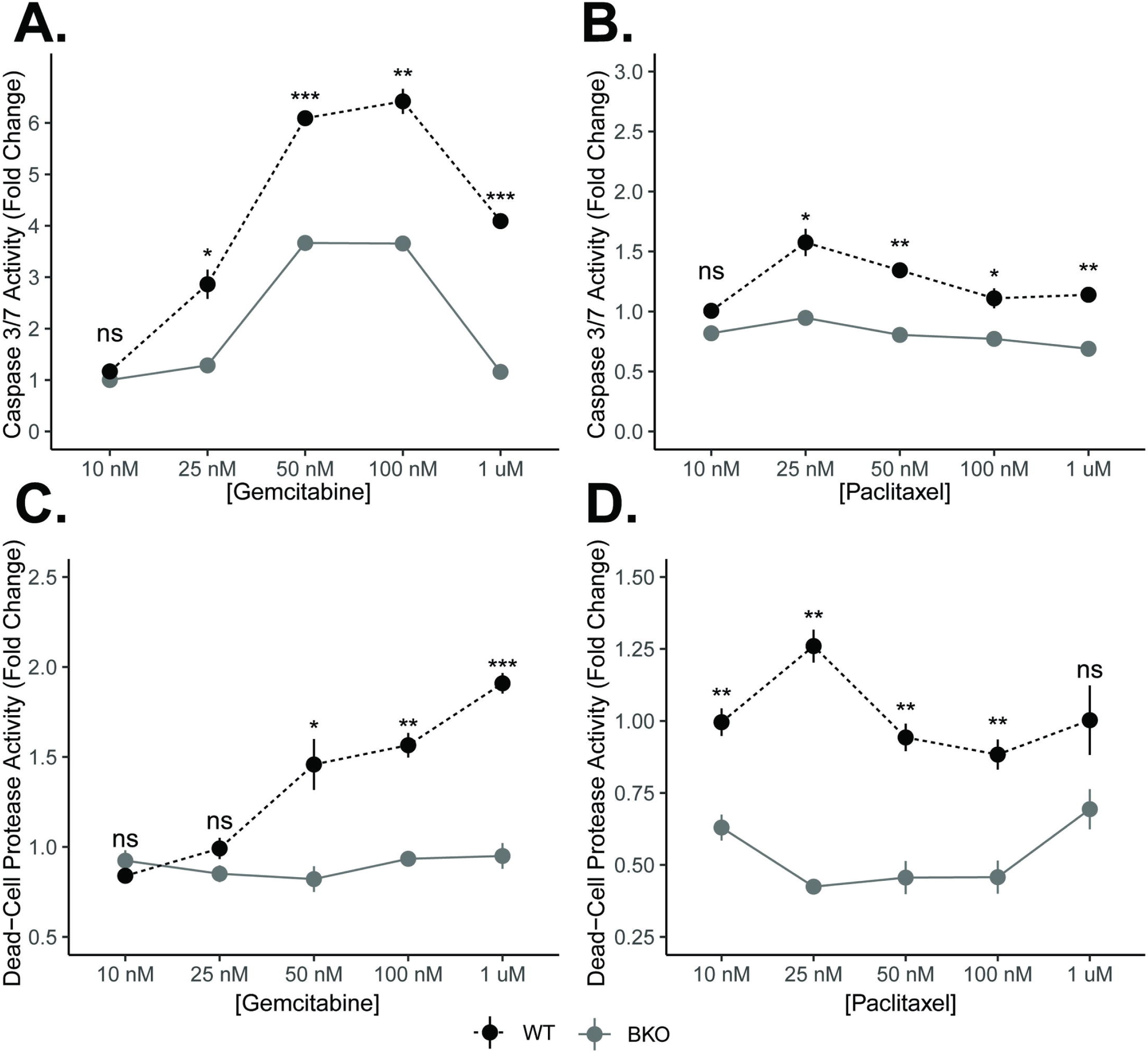
Clock dysfunction diminishes the chemotherapeutic cell death response. KPC wildtype (WT) and KPC-BKO cells (n = 3) were treated with increasing doses of gemcitabine and paclitaxel. Fold change differences (± standard error) in Caspase 3/7 activity in response to either **A.** gemcitabine or **B.** paclitaxel. Fold change differences (± standard error) in dead-cell protease activity in response to **C.** gemcitabine or **D.** paclitaxel. Comparisons between conditions at each concentration were made with t-test. [ns = not significant, * = *p* < 0.05, ** = *p* < 0.01, *** = *p* < 0.001]

**Fig 9:**
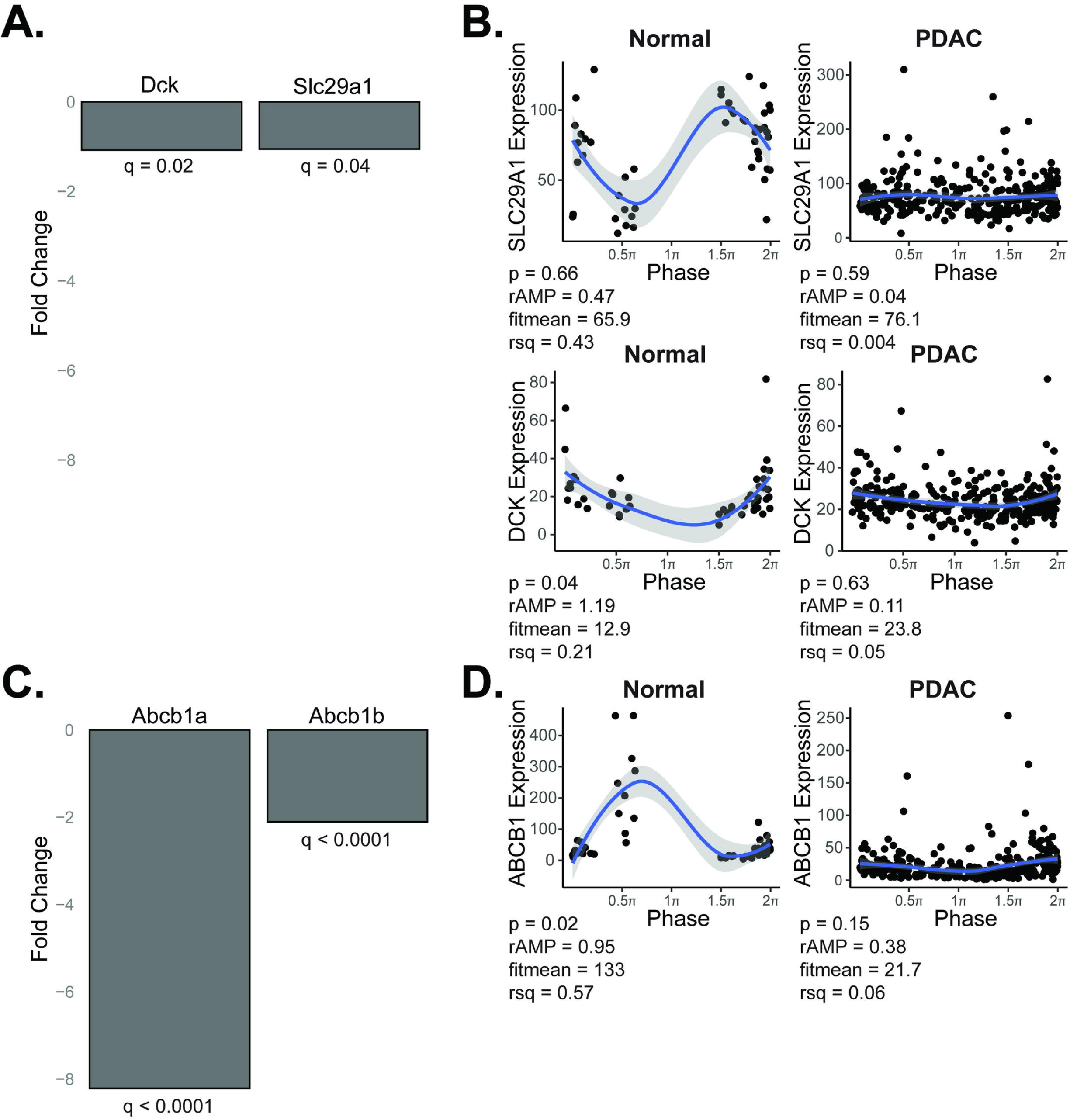
Evaluating for mechanisms of chemotherapy resistance due to clock dysfunction. <u>Gemcitabine resistance mechanism</u>: **A**. Fold change in expression between KPC-BKO and KPC for channel protein hENT1 (*SLC29A1* gene) and deoxycytidine kinase (*DCK* gene) is shown, with negative values representing downregulation in the KPC-BKO (clock disrupted) cells. **B.** CYCLOPS reordered samples (0 to 2π) from normal (n = 50) and pancreatic ductal adenocarcinoma (n = 318) is depicted for each of the respective genes from (**A**), demonstrating arrhythmic gene expression in human normal and pancreas cancer tissue. Shading around the blue regression line indicates the 95% confidence interval. <u>Paclitaxel resistance mechanism</u>: **C.** Fold change in expression between KPC-BKO and KPC for ATP-binding cassette drug efflux transporters (*Abcb1a and Abcb1b* genes) are shown, with negative values representing downregulation in the KPC-BKO (clock disrupted) cells. **D.** CYCLOPS reordered samples (0 to 2π) depict the human *ABCB1* gene expression in normal (rhythmic) and PDAC (arrhythmic and suppressed) tissue. Shading around the blue regression line indicates the 95% confidence interval. *p* < 0.05, rAMP > 0.1, goodness of fit (rsq) > 0.1, and fitmean > 16 were considered rhythmic for CYCLOPS analysis (27).

## Discussion

Our work herein used matched normal and tumor samples from TCGA and CPTAC-3 to provide the first substantive evidence, to our knowledge, that the circadian clock is disrupted in PDAC while the adjacent normal pancreatic clock is intact. Recently published work by Talamanca *et al.* also demonstrated an intact circadian clock in human pancreas, albeit pancreas from human donors and not tumor-adjacent pancreas (51). The authors leveraged transcriptional data from the GTEx consortium and used an algorithm to project phases across tissues in postmortem human donors based on a donor internal phase. They presumed that relative circadian phases are consistent among individuals to identify sex- and age- dependent changes in circadian expression. Although the main metabolic organ of focus was the liver, in the pancreas they identified a relatively strong concordance of rhythmically expressed genes between males and females. However, when comparing pancreatic tissue from young versus old individuals, there was a significant decrease in the percentage of conserved rhythmically expressed genes. Meanwhile, Wucher et al also used the GTEx consortium data and time of death (as well as the season of death) to determine differentially expressed genes between the day and night and across seasons (52). With this method, they were able to identify differentially expressed genes between the day and night across tissues, and they found that among circadian clock genes, *PER1-3* demonstrated the lowest logFC (night/day) while *BMAL1* and *NPAS2* showed the highest logFC (night/day). Interestingly, *BMAL1* was the single gene with a day-night pattern across the greatest tissue number (33 tissues). Also of note, the authors identified a seasonal variation in gene expression with approximately 5% of the pancreatic transcriptome demonstrating differential expression in at least one season when compared to the other seasons. These results have implications when evaluating population-level data and indicate sex, age, and potentially the time of year are important factors depending on the tissue assessed. To evaluate clock functionality in the pancreas in our study, we examined the clock with three complementary analyses not previously performed in pancreatic tissues: nCV, clock correlation, and CYCLOPS. The nCV was pioneered and validated in several tissues by Wu and colleagues (25). It is a measurement that is directly related to the relative amplitude (rAMP) of the oscillation and assesses overall and individual clock gene robustness (25). For example, Wu and colleagues demonstrated that the nCV of core clock genes (CCGs) was consistently and significantly diminished in clock-disrupted tissues versus wild-type tissues (*e.g*., *Bmal1* knockout adipocytes vs wild-type adipocytes) and human datasets of tumor compared to matched (adjacent) normal tissue, where the timing of sample collection was unspecified (25). It is therefore extraordinarily beneficial for understanding the rAMP of the circadian clock genes – a key measure of clock health – in population-level data where sample time acquisition is unknown. In conjunction with nCV, Shilts *et al*. developed the method of clock correlation to determine the progression of the clock in time-indeterminate datasets (22). By capitalizing on the concept of co-expression of CCGs (intrinsic to the transcriptional translational feedback loop), Shilts and colleagues were able to computationally discern clock progression in transcriptomic data from numerous human datasets. Importantly, this technique is not dependent on full coverage of the period by samples, providing a beneficial approach for our normal data set which included only 50 samples and did not have phase representation across the 24- hour period. Furthermore, a direct comparison between heatmaps can be made (*i.e*., murine vs human or tumor vs normal) because each heatmap has the same corresponding color to rho correlation value (22). Thus, when combining nCV and clock correlation analysis in unordered human samples, the health of the circadian clock in population-level data can be ascertained even when working with low sample numbers (at least 30).

Wu and colleagues applied nCV analysis to several paired tumor-normal datasets (25), while Shilts *et al.* examined clock correlation in normal tissues and paired tumor-normal datasets (22). However, we are the first group, to our knowledge, to analyze the pancreas, possibly because our recent publication was the first, to our knowledge, to characterize the diurnal expression of mouse pancreatic genes over 48 hours (23). Thus, an appropriate mouse reference group had not been published for comparison. In the study by Shilts *et al*., the human liver clock correlation heatmap (a similarly metabolic organ) demonstrated a weaker clock correlation versus the mouse reference, which they attributed to increased noise in the liver dataset (22). Therefore, in our human data, we expected to identify an apparent ‘diminished health’ of the clock (lower nCV and weaker clock correlation) in the human pancreatic tissue versus mouse samples due to significant variation in the normal samples. This variability is contributed by human factors, such as type and timing of diet (53,54), underlying genetic differences, and alterations in the ‘normal pancreas’ that surrounds the tumor (pancreatic atrophy, fibrosis, inflammation, etc.) (55). These can modulate the relative amplitude of oscillation or CCG correlation which contributes to significant noise in the data, as compared to the genetically identical, age-matched, and environmentally matched mouse samples (22,28). Regardless of these limitations, the nCV and clock correlation revealed two key components of a healthy clock in human pancreas, which was an important component of the pipeline.

CYCLOPS is a machine learning method developed by Anafi *et al* (28), with subsequent elegant studies by Wu *et al* (26,27)., to demonstrate how clock gene relationships can be used to infer and reorder samples by their predicted phase of expression to understand circadian biology in population-level data. The authors who developed CYCLOPS recommend roughly 250 samples for a complete phase distribution (28), which is dependent on differences in time of surgery (presume specimens acquired between 6 AM and 6 PM) as well as differences in genetics and environmental factors (*e.g*., differences in sleep-wake cycle, or shift worker status) (56). CYCLOPS has been applied to the lung, liver, brain, hepatocellular carcinoma, and skin, but not the pancreas (27,28). This is possibly due to known challenges associated with CYCLOPS, which includes optimizing the seed gene list for appropriate ordering (28). We used the modified CYCLOPS approach by Wu *et al*. and were able to leverage our murine pancreatic longitudinal expression data to generate the seed gene list for use with human data, which increases the signal-to-noise ratio to optimize ordering capability (23,27,28). Thus, despite the expected variability in human normal pancreas samples, CYCLOPS significantly ordered the samples across the period, with several clock genes, including *CLOCK*, *PER1*, *PER3*, *NR1D2*, *RORC*, *NFIL3,* T*EF*, *NR1D1, BHLHE40,* and *NPAS2* demonstrating rhythmic expression. While *BMAL1* and *CRY1* did not pass our predetermined cutoff for rhythmicity in the normal samples, they demonstrated a robust rAMP with near-significant p values likely reflecting limitations with our sample number (*i.e*., distribution of samples across the period) than true lack of rhythmicity in the normal samples. As evidenced by the apparent gap in phase in the normal data, there was not uniform distribution of normal samples across the 24-hour period, which signifies a limitation of our analysis. Incorporating time-stamped normal samples (the hybrid design) can significantly increase the accuracy of CYCLOPS ordering results, as suggested in Wu et al (26,27). Currently, we cannot implement a hybrid experimental design in this study. However, despite these limitations, CYCLOPS statistically ordered the human normal data, and the phase-set enrichment analysis and predicted phase of expression (of most CCGs) aligned well with our prior mouse pancreas transcriptomic data (reference dataset). When combined with nCV and clock correlation, the data clearly demonstrate an intact clock in the human normal pancreas samples, which was an essential premise for evaluating the clock in human PDAC.

We then proceeded to use nCV, clock correlation, and CYCLOPS to provide compelling evidence of circadian clock disruption in human PDAC. While the concept of circadian disruption in PDAC has been posited by others, past studies examining global clock function in human PDAC have been limited by detecting binary differences in overall RNA and protein expression between normal pancreas and PDAC (17,18,20,21). Relles *et al.* found that several CCGs demonstrated decreased expression in PDAC compared to benign tumors or normal pancreas (18), while Li *et al*. found that low *Bmal1* expression (compared to ‘higher expression’) was associated with worse disease-free and overall survival in patients with PDAC (17). Similar studies have been repeated with concordant findings (20,21). However, binary comparisons of expression are unlikely to capture the complexities of clock health such as rhythmicity, phase changes, changes in rAMP, or changes in clock progression. With the 318 available PDAC samples, we showed that there was markedly diminished nCV and a much weaker correlation among clock genes in PDAC compared to normal pancreas, depicting a loss of clock health in the cancer tissue. Further, although the inability to reorder PDAC samples may be a limitation of CYCLOPS, it more likely indicates clock dysfunction (57). In the PDAC samples, there was a sufficient sample number (n = 318) for phase distribution, but the rAMP of nearly all CCGs was markedly decreased (as visualized through best reordering), with many manifesting an arrhythmic pattern. Collectively, our approach to human data (nCV, clock correlation, and CYCLOPS) provided convincing evidence of clock disruption in PDAC.

The main limitation of our current strategy is the inability to discern why there was a loss of circadian signatures in human PDAC population data. While we have shown compelling evidence from the human sample transcriptomic data that circadian disruption is present in PDAC, the population-level data doesn’t accurately represent individual patient tumors. The literature in cancer would suggest that cohorts of patients exhibit a differential extent of clock disruption in their tumors (*i.e*., some patient tumors have an intact clock while others are markedly disrupted) (15,17,18,39). This is seen in other cancers. For example, in human neuroblastoma, MYC amplification distorts circadian repressors (NR1D1) and circadian activators (RORA and BMAL1) to cause metabolic rewiring that fuels cancer cell growth and promotes poor patient outcomes (15,39). Consequently, clock function in neuroblastoma can be subcategorized based on low/normal MYC expression (intact clock) versus MYC amplification (clock disrupted). Similarly, in human PDAC, the overall loss of circadian signatures seen in the nCV, clock correlation, and CYCLOPS analysis could be the consequence of i) dampening of the clock to a significant extent in every patient (*i.e*., global phenomena), ii) differences in the extent to which the clock is disrupted (*i.e*., clock disrupted vs intact clock), or iii) biological heterogeneity in the PDAC samples surpassing circadian variability. The last consideration seems unlikely given the marked decrease in rAMP (nCV) and clock correlation showing significantly diminished clock health in PDAC. Furthermore, as seen with the CYCLOPS ordered *PER1* expression data over the 24-hr period (**Fig 3A**), enhanced variability did not lead to the apparent arrhythmicity in PDAC. Unfortunately, we are currently unable to discriminate between the first two possibilities and test the hypothesis that patient tumor subgroups harbor differences in clock function. Ideally, we would be able to subcategorize the population of patients instead of evaluating the entire cohort of PDAC patients, given the known molecular heterogeneity in PDAC (58). For example, there are alterations in different gene programs (*e.g.*, *Kras*-mutant, *TGF-β* signaling, G1/S checkpoint signaling, etc.) that determine the molecular subtypes in PDAC such as squamous (including MYC pathway activation), pancreatic progenitor (including programs controlling fatty acid metabolism), immunogenic, and ADEX (58). Therefore, separating the cohort into subtypes to evaluate clock function could help elucidate whether circadian disruption in human PDAC co-segregates with specific molecular subtypes or if there are alterations in specific gene programs that promote clock disruption. However, the strength in our current evaluation derives from the number of patients included in the analysis – and this is requisite for CYCLOPS analysis (minimum of ∼200 patients needed for optimal phase distribution and functioning of the algorithm) (28). Thus, despite batch correction algorithms, the significant heterogeneity in transcriptomic data (*e.g.,* institutional differences in experimental conditions, tumor purity, etc. (59–62)) results in challenges in interpreting differences in clock function and drawing conclusions from smaller subsets of patients, particularly with algorithms like CYCLOPS (28). Therefore, the focus of the current work was to establish that clock dysfunction is present in human PDAC. In future work, we will build on this foundation and integrate whole genome sequencing with RNA sequencing from our institutional samples, combined with uniform pathologic evaluation and consistent sequencing conditions to minimize the variability/batch effect and optimize the signal-to-noise ratio. This strategy will enable us to investigate whether specific molecular subtypes of pancreas cancer co-segregate with clock dysfunction.

Our assessment of clock function in the KPC cells would also suggest that clock function is heterogeneous in PDAC, given that the clock was intact in the *Kras*- and *Trp53*-mutant pancreas cancer cells and necessitated *Bmal1* mutagenesis to generate clock disruption. Further, we assessed the cycling of the *Per1* gene and *Per2*-luciferase in the KPC cells and identified a robust rhythm; by comparison, the human PDAC samples demonstrated arrhythmic *PER1* expression. This discrepancy highlights the challenge associated with the use of immortalized cancer cell lines for extrapolating to circadian clock function and cancer-specific distortion of the clock in human patients. These *in vitro* and *in vivo* models can be used to evaluate the impact of specific clock alterations, which can also yield information on potential novel therapeutic approaches (*e.g.,* uncovering the mechanism through which *Bmal1* knockout causes accelerated tumor growth could lead to therapeutic vulnerabilities). Yet, this type of model system is uniform with respect to underlying PDAC-specific mutations (all lines in our study are derived from the same *Kras* and *Trp53* mutations) and does not recapitulate the unique nature of human PDAC *in situ*, where individual patient tumors exhibit mutational heterogeneity (58). Therefore, we did not have a ‘representative model’ in our immortalized studies of each human PDAC subtype to determine which subtypes may harbor a disrupted clock. Unfortunately, there is currently no methodology to obtain longitudinal data from each individual patient tumor; such an approach would enable an understanding of the circadian clock at the individual level and would help to determine the circumstances under which tumors manifest clock disruption, such as with specific gene program alterations or in the setting of metastatic disease. As noted previously, the strategy of future work to uniformly evaluate subsets of PDAC to determine clock disrupted versus clock intact subtypes can help to address this significant discrepancy.

We generated the novel murine KPC-BKO cell line as a basis to evaluate clock disruption in PDAC. Using our KPC and KPC-BKO cells, we found accelerated tumor growth in our syngeneic *in vivo* model with *Bmal1* functional knockout. Our preference was to use an immune-competent model given the known impact of the circadian clock on the immune system (another strength of our model) (63,64). Interestingly, our primary tumor growth results were similar to results by Jiang and colleagues, who used implanted human PDAC cells (BxPC-3 and PANC-1) into immunocompromised mice after shRNA knockdown of *Bmal1* (20,21). However, patients with PDAC ultimately succumb to distant metastatic spread, rather than local tumor growth, so examining the contribution of clock disruption to overall survival was of greater importance (65). We found that *Bmal1* mutation promoted earlier lethality, which has not been shown before. Concordant with the aggressive tumor phenotype, *Bmal1* mutation also caused resistance to chemotherapy. While resistance to gemcitabine has been shown (21), we found chemoresistance to two different backbone anti-cancer agents (gemcitabine and paclitaxel), including suppressed apoptosis and cytotoxicity, indicating a more broad resistance to standard PDAC chemotherapy. Although suppressed *Bmal1* expression in PDAC has been suggested to modulate *Trp53* to promote a tumor suppressor effect, this was unlikely the case in our study considering KPC cells are a *Trp53*-mutant cell line (20). Other work indicates the transcription factor YY1 is significantly overexpressed in PDAC and ultimately causes BMAL1 suppression with consequent PDAC progression and resistance to gemcitabine (unclear mechanism of resistance) (21). Yet, when we examined our human PDAC and human normal samples, we found that *YY1* expression was equivalent between the groups (mean expression 101.61 versus 102.87). Turning to other cancers where BMAL1 is suppressed, *MYC* amplification in neuroblastoma alters *BMAL1* mRNA expression through the induction of *NR1D1* (39,66). Notably, this is associated with poor prognosis and is *BMAL1* dependent since ectopic expression of *BMAL1* inhibits tumor growth. Given the known interaction between MYC and the circadian clock, and the known presence of MYC pathway activation in PDAC (squamous subtype), this will be a focus in future work to evaluate the molecular subtypes of PDAC (15,39,67–69). However, this mechanism was not apparent at the global population-level since we did not identify upregulation of *MYC*, *MYC-N,* or *NR1D1* gene expression in human PDAC compared to normal samples. While common resistance mechanisms of gemcitabine and paclitaxel were not reflected in the human PDAC versus human normal data or the KPC versus KPC-BKO cells (*e.g*., channel proteins), RNA sequencing identified several enriched pathways integral to cancer progression and chemoresistance in the *Bmal1* functional knockout cells, such as the PI3K-AKT pathway (70,71). The PI3K-AKT pathway is inextricably linked to cancer cell proliferation and resistance to apoptosis, indicating a plausible mechanism for inhibition of programmed cell death to multiple agents seen in our study (72,73). Further, resistance to paclitaxel is associated with the activation of the PI3K-AKT pathway (74), and similar correlations have been identified with gemcitabine resistance (75,76). However, the mechanism of chemoresistance is quite complex and the etiology for suppressed apoptosis and cytotoxicity in the KPC-BKO line versus the KPC line remains unclear. To add additional complexity (and highlight another limitation), it is currently unresolved whether the effects we have seen with the loss of functional BMAL1 protein are due to clock disruption or an independent role. Furthermore, clock-dependent effects can depend on which arm is perturbed, the positive arm (circadian activators) or negative arm (circadian repressors) of the clock. For example, in recent seminal work by Liu and colleagues, abolishing different arms of the circadian clock resulted in reciprocal effects on the expression of *c-Myc* and downstream target genes (68), with the knockout of *Bmal1* causing increased expression and *Cry1/2* knockout causing suppressed expression of *c-Myc* and *c-Myc* regulated genes, respectively. Therefore, in future work, we can employ a series of clock-manipulated PDAC cell lines (*e.g., Per1/2* double knockout to disrupt the negative arm) to better understand the contributions of clock disruption and various components of the clock to chemoresistance and PDAC progression.

In conclusion, we used a comprehensive approach (nCV, clock correlation, and CYCLOPS) to evaluate the health of the circadian clock in human normal pancreas and demonstrated clock disruption in human PDAC. Additionally, we developed novel cell lines to evaluate the repercussions of clock disruption in PDAC and identified factors associated with poor prognosis (*i.e*., worse survival, resistance to chemotherapy, and enrichment of cancer-related pathways). While we acknowledge that significant work needs to be done to improve our understanding of the circadian clock in PDAC, we can use the foundation from this study as a basis for future work, where we will disentangle the clock-dependent effects of PDAC and consequently focus therapeutic efforts.

## Materials and Methods

### Mouse Care

All animal studies were conducted according to an approved protocol (M005959) by the University of Wisconsin School of Medicine and Public Health (UW SMPH) Institutional Animal Care and Use Committee (IACUC). Male and female C57BL/6J mice were housed in an Assessment and Accreditation of Laboratory Animal Care (AALAC) accredited selective pathogen-free facility (UW Medical Sciences Center) on corncob bedding with chow diet (Mouse diet 9F 5020; PMI Nutrition International) and water ad libitum. All *in vivo* experiments were performed with age-, cage-, and sex-matched mice under identical lighting conditions. All experiments were performed with mice 6-8 weeks of age, and cages were selected at random for each experiment.

### Clock Correlation, nCV, and CYCLOPS Pipeline

We assessed the overall clock gene correlation and robustness of the clock with the clock correlation matrix and normalized coefficient of variation (nCV) – nCV is known to be correlated with the relative amplitude (rAMP) of oscillating clock genes, indicating the clock robustness as previously described (22,25). The nCV was calculated for the overall condition with the nCVnet and nCVgene functions (25). Clock correlation matrices were created using an available shiny app (https://github.com/gangwug/CCMapp) which compares the correlation of clock components (17 individual clock genes) to a baseline correlation from the circadian atlas using the Mantel test (22,26,27,31,32).

Cyclic ordering by periodic structure (CYCLOPS) (28) was validated for use in the pancreas utilizing our existing murine normal circadian and chronic jetlag pancreas RNA sequencing (RNA-seq) data (Gene Expression Omnibus (GEO) Accession number: GSE165198) (23). Specifically, the seed genes for use in CYCLOPS were selected by cross-referencing genes found to be rhythmically expressed in our dataset with those genes either rhythmically expressed in the liver (similarly metabolic organ) or those used by Wu *et al.* when validating CYCLOPS in the skin (**Supplemental Data File 3**) (23,26,31,32). The updated CYCLOPS pipeline by Wu *et al*. (https://github.com/gangwug/Oslops) was then used to reorder our murine pancreas datasets with known sample collection times (26). Eigengenes were selected with the Oscope package to sharpen CYCLOPS (77). Clusters with a *p* < 0.05 (also known as Stat^err^) and Met^smooth^ < 1 were considered to be significantly reordered (28). Rhythmicity of the reordered genes was determined on cosinor analysis with a *p* < 0.01, rAMP > 0.1, goodness of fit (rsq) > 0.1, and fitmean > 16 (26). Significantly rhythmic gene phase was then compared to the rhythmic gene phase detected from the known sample time collection using the meta3d function of Metacyle (42). Clock genes were highlighted to demonstrate a correlation between the predicted and actual phase.

The clock was evaluated in human normal and human PDAC RNA-seq datasets from TCGA and CPTAC-3 (29,30). After batch correction with ComBat and filtering, 50 matched normal and 318 PDAC samples were obtained for analysis (78). The pipeline described above was then used to obtain the clock correlation matrix, nCV, and CYCLOPS reordering. Cosinor analysis was performed to test for rhythmicity. Given the additional biologic heterogeneity of the human data, a *p* < 0.05, rAMP > 0.1, goodness of fit (rsq) > 0.1, and fitmean > 16 were used as a rhythmicity cutoff. Rhythmic genes from normal samples were assessed with phase set enrichment analysis (PSEA) (79). Rhythmic gene sets ordered by significance were inputted with their calculated phase of expression. Default settings were used for PSEA, including domain 0-24, min item 10, max sims/test 10,000. The gene set enrichment analysis (GSEA) gene ontology (GO) (c5.go.bp.v7.5.1.symbols) set was leveraged as the pathway input. The top 15 significant pathways (*q* < 0.05) were selected for representation.

### KPC Cell Line Creation and Maintenance

Pancreas cancer cells that harbor *Kras^G12D^* and *Trp53^R172H^*mutations (KPC cells) were acquired from Ximbio (Catalog: 153474; Westfield Stratford City, UK). Cells were cultured in DMEM supplemented with 10% Fetal Bovine Serum (Cytiva, Marlborough, MA), 1% L-glutamate-Penicillin-Streptomycin (Gibco, ThermoFisher Scientific, Waltham, MA), and 1% non-essential amino acids (Gibco) at 37°C at 5.0% CO_2_ to their appropriate confluence for use. CRISPR/Cas9 technology was used to introduce *Bmal1* mutations into the well-established KPC cells, generating the *Bmal1* functional knockout line (KPC- BKO) (36). Synthetic tracrRNA and target-specific crRNAs (crRNA:tracrRNA (ctRNAs)) were annealed as per manufacturer instructions (Integrated DNA Technologies (IDT), Coralville, IA).

Ribonucleoproteins were formed with Cas9 protein (V3, Catalog: 1081059, IDT) and ctRNAs individually (ctRNA 1:AATATGCAGAACACCAAGGA, ctRNA 2: TTAGAATATGCAGAACACCA) (IDT). Nucleofection was used to introduce RNPs (1.95 µM Cas9, 2 µM ctRNA, 2 µM electroporation enhancer (IDT)) into 2 x 10^5^ KPC cells on a Nucleofector 4D (Lonza Biosciences, Walkersville, MD) with an SF kit (Catalog: V4CX-2032, Lonza Biosciences) as per manufacturer instructions using pulse code CM-120. After 48 hours of recovery in DMEM growth media (described above) at 37°C at 5.0% CO_2_, single cells were deposited in 96 well plates using a BD FACSAria III (BD Biosciences, Franklin Lakes, NJ). Outgrowing clones were condensed to a 96 well plate in duplicate to propagate clones and generate a genomic DNA source. Genomic DNA was harvested from KPC cells, nucleofected with ctRNA-1 RNPs and ctRNA-2 RNPs, and the targeted region of *Bmal1* was PCR amplified using primers: Forward: acactctttccctacacgacgctcttccgatct NNNNNN CCAAGAATCCTTGTGTGTCTG and Reverse: gtgactggagttcagacgtgtgctcttccgatct AGAGGACTCCACAGACATGAAC (IDT). PCR products were dual-indexed with indexing PCR, pooled, sequenced on an Illumina MiSeq instrument (San Diego, CA), and analyzed with CRISPResso2 (80).

### Creation of Per2-dLuciferase reporter KPC cell line

KPC cells were stably transfected with a mammalian gene expression vector harboring a destabilized luciferase reporter driven by the *Per2* promoter fused to intron 2 of *Per2* and a puromycin resistance cassette (VectorBuilder, Santa Clara, CA) (81). The vector was transfected with lipofectamine 2000 (ThermoFisher Scientific) and incubated for 2 days, and then exposed to media containing 2.5 μg/mL puromycin for 3 days, and surviving clones were selected. Luciferase activity was measured in the selected clones with the luciferase assay system (Promega, Madison, WI) on a BMG CLARIOstar (BMG Labtech, Ortenberg, Germany) plate reader. The two selected clones were then subcloned using a BD FACSAria III after staining with DAPI. Luciferase activity was again measured in the subclones to validate expression before use in downstream experiments.

### Clock Function Testing

To evaluate for clock function, KPC and KPC-BKO cells were synchronized with 200 nM dexamethasone for 2 hours in FBS-free DMEM media, followed by RNA isolation 24 hours after the synchronization using the RNeasy protocol (Qiagen, Hilden, Germany) according to the manufacturer’s recommendations (82). Samples (n = 3 biologic replicates) were collected at 4-hour intervals for 24 hours. Quantitative real-time polymerase chain reaction (RT-qPCR) was performed (in technical triplicate) for the downstream core clock gene (CCG) *Per1* (ID: Mm00501813_m1, Life Technologies, Carlsbad, CA) and the housekeeper gene *Hprt* (ID: Mm03024075_m1) using GoTaq Probe qPCR and RT-qPCR System (Promega, Madison, WI) and Quantstudio 7 flex RT-PCR system (ThermoFisher Scientific). Expression was measured and the ΔCT was calculated. The mean ΔCT values were then tested for rhythmicity using the meta2d function of Metacycle (42). For Metacycle settings, the min period and max period were set to 24, and “JTK”, “LS”, and “ARS” were selected for cycMethod. An integrated FDR corrected *q* value < 0.05 and rAMP > 0.1 were taken as rhythmic as previously described (26). To separately evaluate clock function in KPC cells, luciferase activity driven by the *Per2* promoter was measured. Cells from two independent clones were synchronized with 200 nM dexamethasone for 2 hours in FBS-free DMEM media, and 24 hours later luciferase activity was measured at 4-hour intervals (n = 6 technical replicates per time point). To measure activity, 1x cell culture lysis reagent (Promega) was added and cells were incubated for 5 minutes. Firefly luciferase assay reagent (Promega) was added and luminescence was measured on a BMG CLARIOstar plate reader and tested for rhythmicity using Metacycle.

### Western Blotting

Western blotting was performed to determine BMAL1 expression in two independent KPC cell lines. After synchronization, protein samples were isolated at 6-hour intervals for 24 hours using CelLytic M lysis reagent (MilliporeSigma, Burlington, MA) and Halt Protease and Phosphatase Inhibitor Cocktail (ThermoFisher Scientific). A total of 30 μg of each sample was loaded onto a Mini-PROTEAN TGX 7.5% precast mini-gel (Bio-RAD Laboratories, Hercules, CA). The gel was then transferred using the semi-dry transfer technique to an Immobilon-P PVDF membrane (MilliporeSigma). The membrane was then blocked with 5% skim milk and incubated with 1 μg/μL rabbit BMAL1 antibody (Catalog: NB100- 2288, RRID: AB_10000794, Novus Biologicals, Littleton, CO) and 1:2000 rabbit β-ACTIN antibody (Catalog: 4967, RRID: AB_330288, Cell Signaling Technologies, Danvers, MA). Finally, the membranes were incubated with alkaline phosphatase-conjugated goat anti-rabbit IgG (1:10000) (Catalog: 111-055- 144, RRID: AB_2337953, Jackson ImmunoResearch West Grove, PA) and stained using 1-Step NBT/BCIP solution (ThermoFisher).

### RNA Isolation, Sequencing, Differential Gene Expression Analysis

To evaluate for transcriptomic differences between wild-type and BKO KPC cells, bulk RNA-seq was performed on 6 independent samples collected from each condition. Isolation was carried out as above and quality was tested for an RNA integrity number (RIN) > 7.5 on the Agilent 2100 bioanalyzer (Agilent Technologies, Santa Clara, CA). A total of 300 ng of mRNA was enriched with poly-A selection and sequencing on the Illumina HiSeq2500 platform by the University of Wisconsin Biotechnology Sequencing Core. FASTq files were processed with Skewer and genes were filtered to remove those with low expression (83). Samples were normalized by the method of trimmed mean of M-values (84). Contrasts were drawn with the edgeR package, with differential expression taken when the FDR *q* < 0.05 (45). Pathway testing was performed with the KEGG database (Kyoto Encyclopedia of Genes and Genomes) using previously described methods (85). The top 500 significant genes were inputted, ordered by *q* value, and the top 9 significant pathways (where *p* < 0.05) were plotted for visualization. Pathway dot size is indicative of the number of genes in each pathway. The RNA-seq data is publicly available through GEO (Accession number: GSE213680).

### Heterotopic Tumor Modeling

To create flank tumors, 1 x 10^5^ KPC or KPC-BKO cells were injected into the right flanks of immunocompetent C57BL/6J mice obtained from Jackson Laboratory (RRID: IMSR_JAX:000664, Bar Harbor, ME). Cells were mixed in a 1:1 50 μL solution of DMEM media and Matrigel (Corning Inc, Corning, NY). In the tumor growth experiment, a single dose of KPC or KPC-BKO cells was injected into C57BL/6J mice (male: n = 5, female: n = 5, each group) and tumors were measured twice weekly for four weeks starting on day 6 with the caliper method as previously described (86). Tumor length and width were measured and tumor volume was calculated using the formula: tumor volume = length x width^2^ x ½. Tumor weight was also measured (in mg) at the conclusion of the study period. Two independent replicate experiments were performed for tumors derived from KPC cells and KPC-BKO cells (total n = 20 mice per group). Additionally, two separate KPC-BKO clones were tested. Our power analysis, based on prior literature, indicated that using 10 mice per group would have 80% power to detect a 1.5-fold difference (alpha = 0.05) in tumor size (21 days after heterotopic injection) between KPC and KPC-BKO cells (87). Mean differences between groups were computed using a t-test. In the tumor metastasis/survival experiment, the same dose of KPC or KPC-BKO cells was injected into C57BL/6J mice (n = 7 in each group) and tumors were measured weekly until the mice became moribund or died. Kaplan Meier log-rank analysis was then performed to compare survival differences between conditions with the survival package (88). A *p <* 0.05 is taken as significant for both experiments.

### Cell cycle analysis

To evaluate the cell cycle of KPC and KPC-BKO cells, cells were grown to confluence, fixed with 70% ethanol, and stained with 50 μg/mL propidium iodide (Catalog: 421301, BioLegend, San Diego, CA) and 100 μg/mL RNAse A (Catalog: FEREN0531, Thermo Fisher). Samples (n = 3 technical replicates, each condition) were sorted with an Attune NxT flow cytometer (Thermo Fisher) and the DNA content was analyzed for the percent of cells in G1, S, and G2 phase with ModFit LT 6.0 software at the University of Wisconsin Carbone Cancer Center Flow Cytometry Laboratory. The average coefficient of variation (CV) of each sample was < 6%. Differences between conditions were tested with a t-test and a *p* < 0.05 was taken as statistically significant. This experiment was performed independently twice.

### Chemotherapeutic Resistance Evaluation

To evaluate for chemotherapeutic resistance, an equal number of KPC or KPC-BKO cells (3000 each) were grown in 96-well plates for 24 hours. Cells were washed and media containing either vehicle (DMSO, Catalog: D2650-100ML, MilliporeSigma) or increasing concentrations of gemcitabine (dose range = 10 nM to 1μM) (Catalog: G6423-50MG, MilliporeSigma) or paclitaxel (dose range = 10 nM to 1μM) (Catalog: AAJ62734MC, Thermo Fisher) was added to each well. Apoptosis was measured by detecting Caspase 3/7 activity after 24 hours of treatment using the Caspase-Glo 3/7 Assay System (Promega) on a BMG CLARIOstar plate reader. Dead-cell protease activity was similarly measured in cells after 20 hours of treatment using the CytoTox-Glo Cytotoxicity Assay (Promega) on a BMG CLARIOstar plate reader. Mean differences in the fold change between chemotherapy- and vehicle-treated cells were made with a Student’s t-test and a *p* < 0.05 was taken as statistically significant. Each experiment was completed with technical triplicates and independently replicated twice. A representative result is shown for each experiment.

### Analysis

All analyses were performed in R version 4.2.0 or Julia version 0.3.12 unless otherwise indicated. Samples were not excluded from the analysis unless explicitly stated. Standard comparisons of the mean were performed with a t-test and survival was assessed with the log-rank test unless explicitly stated.

## Acknowledgments

We would like to acknowledge the Gene Expression Center and the Bioinformatics Resource Center (BRC) at the University of Wisconsin, Madison for their contributions to this work. The authors thank the University of Wisconsin Carbone Cancer Center Flow Cytometry Laboratory, supported by P30 CA014520, for use of its facilities and services. We would also like to thank the Michael W. Oglesby Foundation for their funding support of our work in circadian disruption and pancreas pathology.

## Materials Availability Statement

The modified KPC cell lines (KPC-BKO and KPC Per2-dLuc) created for use in this manuscript are available upon reasonable request from the corresponding author, Sean Ronnekleiv-Kelly.

## Data Availability Statement

The TCGA and CPTAC-3 data were requested from the following URL: https://portal.gdc.cancer.gov/. All RNAseq data can be found on the Gene Expression Omnibus (GEO) at accession numbers GSE213680 and GSE165198.

## Supplementary Files

### Supplementary Figures

**Supplemental Fig 1:** *CYCLOPS accurately reorders wildtype mouse pancreas samples*. Eigengenes identified by Oscope for use in CYCLOPS in **A.** normal circadian (upper) and chronic jetlag (lower) samples. Shading around the blue regression line indicates the 95% confidence interval for each plot. **B.** The normal circadian (left) and chronic jetlag (right) clusters ordered by CYCLOPS demonstrate accurate reordering for both conditions **C.** Normal circadian (left) and chronic jetlag (right) genes found to be significantly rhythmic on both CYCLOPS reordered cosinor analysis and rhythmicity testing based on the known sample collection time with the Metacycle meta3d function are ordered by their predicted phase of expression. Clock genes are shown in orange.

**Supplemental Fig 2:** *Normal pancreatic and pancreatic ductal adenocarcinoma (PDAC) samples were processed and filtered*. TCGA and CPTAC-3 samples were processed and batched corrected. An MDS plot is shown demonstrating differences between the matched normal (green; n = 50) and PDAC (purple; n = 318) samples.

**Supplemental Fig 3:** *KPC cells expressing dLuciferase driven by the Per2 promoter exhibit circadian activity*. Two independent clones were examined, and luciferase activity was measured at 4-hour intervals (n = 6 per time point). Rhythmicity was calculated with Metacycle. Clone 1 (left) was found to be rhythmic with a *q* = 5.51E-5 and rAMP = 0.44. Clone 2 (right) was found to be rhythmic with a *q* = 1.95E-7, rAMP = 0.39.

**Supplemental Fig 4:** *Cross-validation of mutant Bmal1 (functional knockout) growth in a second independent clone*. A second independent *Bmal1* knockout (BKO) clone with an identical mutation to the first was validated with **A.** western blot analysis. We then heterotopically implanted wildtype (WT) and BKO cells into C57BL/6J mice and followed **B.** tumor weight and **C.** cell growth. BKO_2 (dark grey; n = 20) had a similar mean (± standard error) tumor weight to BKO_1 (light grey; n = 20) (426.09 (± 40.07) mg vs 438.02 (± 48.84) mg; p = 0.85), and both BKO_2 (426.09 (± 40.07) mg vs 280.11 (± 42.73) mg; p = 0.017) and BKO_1 (438.02 (± 48.84) mg vs 280.11 (± 42.73) mg; p = 0.02) were larger than WT (white; n = 20). BKO_1 and BKO_2 had similar significantly faster growth trajectories. **D-G.** KPC wildtype (WT) and BKO_2 cells (n = 3) were treated with increasing doses of gemcitabine and paclitaxel. Fold change differences (± standard error) in Caspase 3/7 activity in response to either **D.** gemcitabine or **E.** paclitaxel. Fold change differences (± standard error) in dead-cell protease activity in response to **F.** gemcitabine or **G.** paclitaxel. Comparisons between conditions at each concentration were made with t-test. [ns = not significant, * = p < 0.05, ** = p < 0.01, *** = p < 0.001]

### Supplementary Data Files

**Supplemental Data File 1:** Significant genes on cosinor analysis for CYCLOPS reordered normal pancreas

**Supplemental Data File 2:** KPC wildtype versus BKO edgeR differential gene expression

**Supplemental Data File 3:** CYCLOPS seed gene list.

**Supplemental Data File 4**: Data for Reconstruction of Figures ***Supplementary Tables***

**Supplemental Table 1: Cosinor rhythmicity analysis of CYCLOPS ordered normal pancreas and PDAC**

