## Supplementary figures and images for "The circadian clock is disrupted in pancreatic cancer"

### Supplemental Figure 1

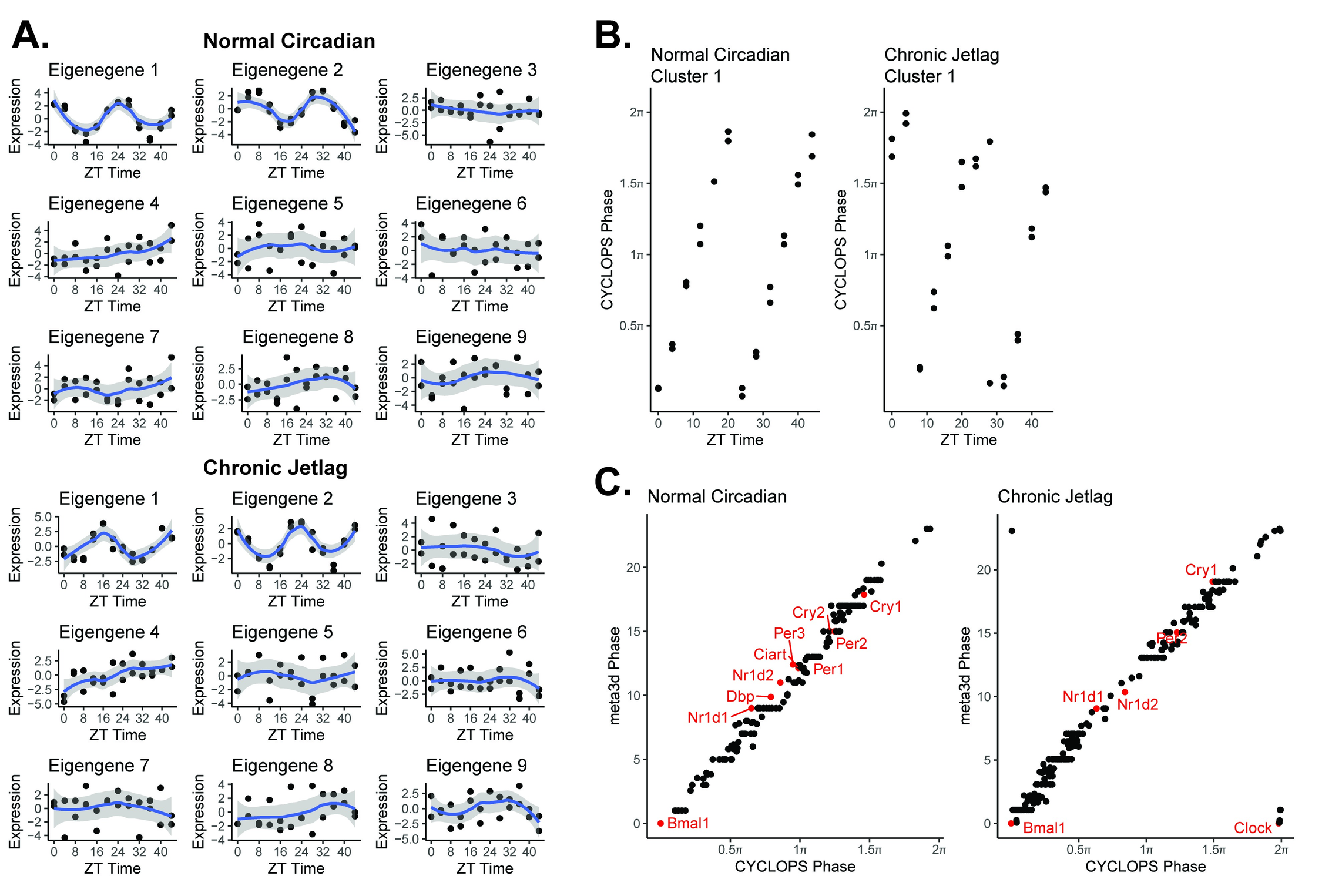

### Supplemental Figure 2

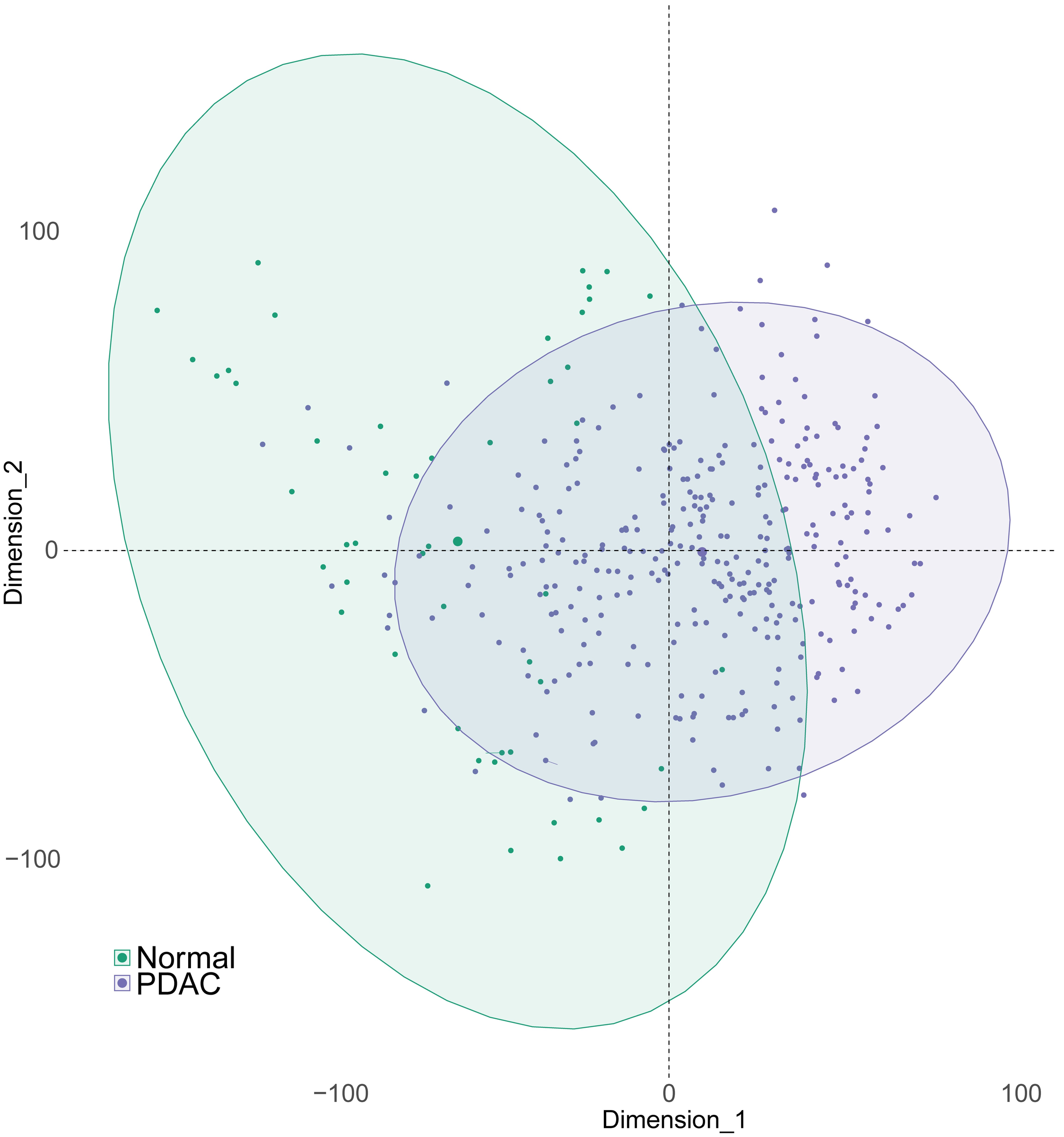

### Supplemental Figure 3

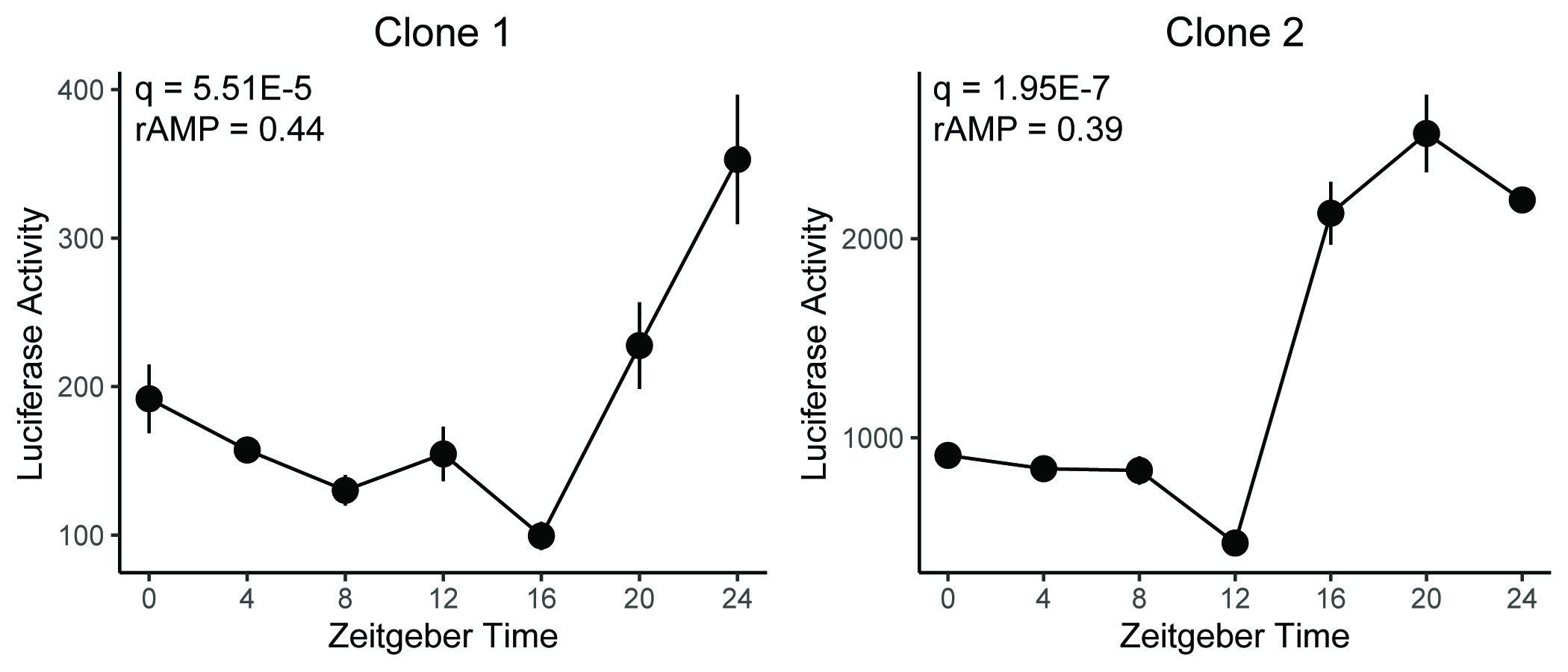

### Supplemental Figure 4

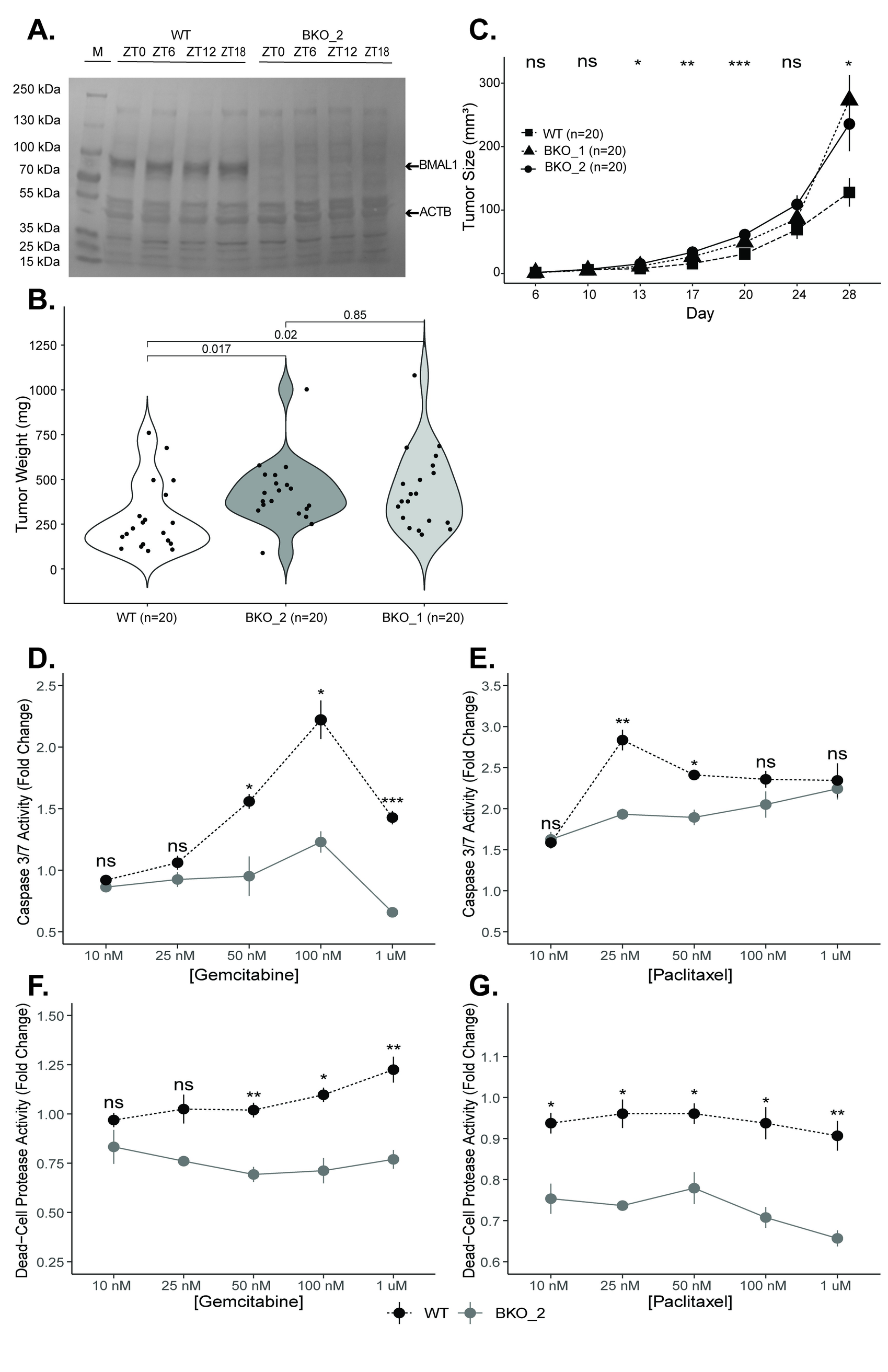
