## Supplemental Table 1 for "The circadian clock is disrupted in pancreatic cancer"

| **Supplemental Table 1: Cosinor rhythmicity analysis of CYCLOPS ordered normal pancreas and PDAC** | | | | | |
| --- | --- | --- | --- | --- | --- |
| **Gene** | **Condition** | **p value** | **rsq** | **fitmean** | **rAMP** |
| *BMAL1* | Normal | 0.071 | 0.245 | 11.674 | 0.796 |
|  | PDAC | 0.31 | 0.06 | 14.654 | 0.135 |
| *CLOCK* | Normal | 0.041 | 0.144 | 55.143 | 0.137 |
|  | PDAC | 0.964 | 0.025 | 58.016 | 0.065 |
| *PER1* | Normal | 0 | 0.363 | 262.898 | 0.805 |
|  | PDAC | 0.453 | 0.001 | 70.816 | 0.025 |
| *PER2* | Normal | 0.198 | 0.252 | 59.603 | 0.475 |
|  | PDAC | 0.724 | 0.011 | 34.16 | 0.048 |
| *PER3* | Normal | 0 | 0.647 | 113.098 | 0.935 |
|  | PDAC | 0.668 | 0.031 | 41.559 | 0.128 |
| *CRY1* | Normal | 0.084 | 0.324 | 24.185 | 0.48 |
|  | PDAC | 0.382 | 0.036 | 22.995 | 0.095 |
| *CRY2* | Normal | 0.159 | 0.099 | 85.107 | 0.274 |
|  | PDAC | 0.995 | 0.007 | 54.098 | 0.065 |
| *NR1D1* | Normal | 0.001 | 0.591 | 35.142 | 1.088 |
|  | PDAC | 0 | 0.299 | 20.914 | 0.481 |
| *NR1D2* | Normal | 0.001 | 0.495 | 89.438 | 0.66 |
|  | PDAC | 0.519 | 0.006 | 57.856 | 0.044 |
| *RORA* | Normal | 0.075 | 0.362 | 41.354 | 0.738 |
|  | PDAC | 0 | 0.234 | 45.003 | 0.289 |
| *RORB* | Normal | 0.089 | 0.234 | 0.799 | 3.855 |
|  | PDAC | 0 | 0.186 | 2.27 | 0.547 |
| *RORC* | Normal | 0 | 0.566 | 74.879 | 0.827 |
|  | PDAC | 0.742 | 0.012 | 24.186 | 0.102 |
| *NPAS2* | Normal | 0.027 | 0.428 | 63.313 | 0.684 |
|  | PDAC | 0 | 0.221 | 40.489 | 0.274 |
| *NFIL3* | Normal | 0.009 | 0.35 | 23.965 | 1.168 |
|  | PDAC | 0.101 | 0.021 | 34.036 | 0.101 |
| *BHLHE40* | Normal | 0.037 | 0.286 | 76.165 | 1.915 |
|  | PDAC | 0.001 | 0.11 | 370.951 | 0.241 |
| *BHLHE41* | Normal | 0.46 | 0.439 | 19.814 | 1.065 |
|  | PDAC | 0.847 | 0.002 | 46.018 | 0.038 |
| *DBP* | Normal | 0.13 | 0.452 | 29.03 | 0.769 |
|  | PDAC | 0 | 0.103 | 12.984 | 0.394 |
| *CIART* | Normal | 0.001 | 0.531 | 8.808 | 0.758 |
|  | PDAC | 0 | 0.193 | 5.953 | 0.441 |
| *TEF* | Normal | 0 | 0.774 | 87.964 | 0.922 |
|  | PDAC | 0.395 | 0.004 | 28.167 | 0.048 |
| *HLF* | Normal | 0.954 | 0.038 | 16.53 | 0.321 |
|  | PDAC | 0.132 | 0.095 | 7.421 | 0.447 |
| *SLC29A1* | Normal | 0.657 | 0.434 | 65.913 | 0.471 |
|  | PDAC | 0.586 | 0.004 | 76.079 | 0.041 |
| *DCK* | Normal | 0.045 | 0.214 | 12.925 | 1.194 |
|  | PDAC | 0.626 | 0.050 | 23.812 | 0.112 |
| *ABCB1* | Normal | 0.015 | 0.573 | 133.049 | 0.950 |
|  | PDAC | 0.150 | 0.057 | 21.682 | 0.383 |
| PDAC - Pancreatic ductal adenocarcinoma, rsq - the goodness of fit, rAMP - relative amplitude | | | | | |
